# Cell cycle expression heterogeneity predicts degree of differentiation

**DOI:** 10.1101/2024.07.19.604184

**Authors:** Kathleen Noller, Patrick Cahan

## Abstract

Methods that predict fate potential or degree of differentiation from transcriptomic data have identified rare progenitor populations and uncovered developmental regulatory mechanisms. However, some state-of-the-art methods are too computationally burdensome for emerging large-scale data and all methods make inaccurate predictions in certain biological systems. We developed a method in R (stemFinder) that predicts single cell differentiation time based on heterogeneity in cell cycle gene expression. Our method is computationally tractable and is as good as or superior to competitors. As part of our benchmarking, we implemented four different performance metrics to assist potential users in selecting the tool that is most apt for their application. Finally, we explore the relationship between differentiation time and cell fate potential by analyzing a lineage tracing dataset with clonally labelled hematopoietic cells, revealing that metrics of differentiation time are correlated with the number of downstream lineages.

## INTRODUCTION

Fate restriction and cellular differentiation are universal aspects of multi-cellular life. These processes are hallmarks of development, tissue repair and homeostasis [1, 2], and their dysregulation is integral to many diseases such as cancer [3] and premature aging [4]. Inferring the relative positions of cells along fate restriction or differentiation axes would be useful to identify rare [5-9] and heterogeneous [10-12] progenitor populations, to reveal molecular programs that regulate development, and to untangle the dynamic consequences of genetic or other perturbations.

In this paper we refer to the axes of fate restriction and differentiation collectively as “differentiation time”. Methods to computationally assign differentiation time for single cell RNA-seq (scRNA-seq) data have proliferated alongside the rapid increase in scRNA-seq adoption [13]. Differentiation time predictors are distinct from trajectory inference (TI) methods such as Monocle, Slingshot, and Wishbone which require a priori knowledge of start or end points [14-16]. Differentiation time predictors are also an attractive alternative to RNA velocity in cases where the validity of its splicing kinetics assumptions is uncertain [17-20]. Therefore differentiation time predictors, designed specifically to approximate degree of differentiation, aid the selection of start and end points of pseudotemporal trajectories and clarify lineages proposed by TI and RNA velocity.

Many differentiation time predictors are based on cellular properties integral to stem cells, development, and differentiation. For example, SCENT and its successor CCAT estimate the ‘signaling entropy’, or the degree of inconsistency in expression among members of biological pathways as determined by protein-protein interaction networks [21, 22], because uncommitted stem cells exhibit sporadic and uncoordinated expression of lineage-specific programs [23, 24]. CytoTRACE, currently the most accurate differentiation time predictor, is based on the concept that undifferentiated cells transcribe more genes than differentiated cells, irrespective of the degree of expression stochasticity [19].

Here, we describe our method to estimate differentiation time: stemFinder. The stemFinder score is motivated by the correlation of cell cycle gene expression with potency [19, 22], the relationship between cell cycle control and potency [25-32], and the association between gene expression stochasticity and fate potency in stem cells [33, 34]. The relationships between fate potential, differentiation and cell cycle are complex. During development, cell cycle kinetics vary with developmental stage based on the cell population size demands of the nascent organ or tissue [35]. Moreover, during development, cell fate propensity is dependent on stage of cell cycle, as seen with germ-layer-specific cell cycle regulators [36]. Both the overall length of the cell cycle and the relative length of each stage are associated with fate potency and differentiation stage. Finally, while proliferating, cells tend to down-regulate transcriptional programs that define their differentiated form [37]. Given these links between cell cycle, differentiation, and fate restriction, we reasoned that a cell cycle-centric method of differentiation time estimation would be at least as accurate as other methods and may overcome some of their deficiencies.

In brief, stemFinder computes the variability in cell cycle gene expression of a query cell in relation to its nearest neighbors based on distances which exclude cell cycle genes. Our rationale was that such a metric would score highly both groups of cells that are proliferating and thus in a mix of cell cycle stages, and groups of cells entering and exiting quiescence. stemFinder returns dataset-specific single-cell differentiation times ranging from 0 (least differentiated) to a maximal score of 1 (most differentiated). We benchmarked stemFinder along with CytoTRACE and CCAT using 24 UMI-based validation sets spanning multiple sequencing platforms, species, and tissue types. We used four metrics of performance to document distinct features of differentiation time predictors and found that stemFinder outperformed the other methods in three of these metrics each. Finally, by analyzing a lineage tracing dataset with clonally labelled hematopoietic cells [38], we show that differentiation time metrics are correlated with the number of downstream lineages.

## METHODS

### Cell cycle gene expression heterogeneity as a metric of differentiation

Our method, stemFinder, uses Gini impurity to compute a neighborhood-wide variability in cell cycle gene expression as a metric of differentiation time, which encompasses both the fate potential, or number of distinct downstream cell types, able to be generated by a given cell and the extent to which a cell has achieved a specialized form (**Figure 1A**). As shown in **Figure 1B**, stemFinder takes as input a K-nearest neighbors matrix (computed without the influence of cell cycle genes) and a Seurat object containing a log normalized, scaled single-cell gene expression matrix. We subset the gene expression matrix for cell cycle genes, binarize it and iterate through each query cell. stemFinder identifies the nearest neighbors of the query cell then quantifies the probability of the binarized gene expression profile of the neighborhood matching that of the query cell. The Gini index is computed for each gene in the query cell based on this probability as a measure of the neighborhood cell cycle gene expression heterogeneity. The resulting values are summed across all genes and returned as an inverted score ranging from a minimum of zero to a maximum possible score of one, where lower scores correspond to less differentiated cells and vice versa. Please consult **Supplementary Methods** for a thorough description of our pipeline and of the stemFinder algorithm. To quantify stemFinder performance, we compare our predicted differentiation time to assigned ground-truth potency values, representing either the ground truth differentiation time as assigned by known lineage relationships of cell types or the developmental time of cells in an isolated lineage. We quantify performance with four metrics: 1.) the Spearman correlation of single-cell values of ground truth versus differentiation time (“single-cell Spearman correlation”), 2.) the Spearman correlation of the mean ground truth versus quantified potency scores of phenotype-defined clusters (“phenotypic Spearman correlation”), 3.) the percentage recovery of cells with low degree of differentiation or high fate potential (**Figure 1C**), and 4.) the discrimination accuracy in distinguishing the most and least differentiated cells in the dataset (AUC, not illustrated). We also compare the performance of our method to two existing methods, CytoTRACE and CCAT, inverting the scores of both methods to correspond to the directionality of our method and of ground truth differentiation time prior to comparison [19, 22]. Please consult **Supplementary Methods** for a description of our validation datasets, benchmarking strategy, robustness testing, and statistical analyses.

## RESULTS

### Heterogeneity in expression of cell cycle genes is inversely correlated with ground truth differentiation

We first investigate whether heterogeneity of cell cycle gene expression or magnitude of cell cycle gene set expression, as reported in previous studies [19], is the superior surrogate for single-cell potency. We compare the Spearman correlation of single-cell ground truth potency and stemFinder score or cell cycle gene set score, respectively, and we test various cell cycle gene lists as input: S/G2M cell cycle genes (the default input to stemFinder), S/G2M/G1 genes, S genes only, G2M genes only, or G1 genes only. Although both metrics inversely correlate with ground truth potency, cell cycle gene expression heterogeneity is a significantly better correlate of ground truth differentiation than cell cycle gene set score, whether we compare results from all input gene lists **(Supplementary Figure 1A**) or the default stemFinder input gene list (**Supplementary Figure 1B**).

### Multiple comparative metrics enable the contextual application of differentiation time

To provide an example of the accuracy and utility of stemFinder, we analyze the murine dentate gyrus validation dataset sequenced on the 10X platform [39]. The dentate gyrus contains both quiescent and proliferating progenitor cells and is where neurogenesis continues into adulthood. This dataset contains the following cell types, from least to most differentiated: radial glia-like cells (RGL), neuronal intermediate progenitor cells (nIPC), neuroblasts, several immature cell types (the astrocyte, granule cell (GC), pyramidal cell, and oligodendrocyte precursor (OPC)), and the terminally differentiated versions of these lineages (**Figure 2A**). As seen in the murine bone marrow, the least differentiated populations (RGL and nIPC) were comprised predominantly of cells in the G2M and S phases of the cell cycle (**Figure 2B**). Ground truth potency values in this dataset recapitulate differentiation time and are assigned on a phenotypic basis using known lineage relationships described in the original paper [39] (**Figure 2C**).

Although all methods perform well in terms of the correlation with ground truth potency and discrimination accuracy, only stemFinder recognizes a majority of RGL cells as highly potent (Figure 2C-D), with a percent recovery of 63.6% as compared with 30.1% for CCAT and 0.57% for CytoTRACE (CytoTRACE only recovers a single RGL cell). stemFinder selects RGL and nIPC cells as most potent, while CytoTRACE incorrectly selects immature GC and pyramidal cells, neuroblasts, and some OPCs as most potent in addition to some nIPCs and one RGL cell (**Figure 2F**). CCAT also selects nIPCs and RGL cells as most potent, although it assigns a wide range of potency scores to each cell phenotype, making different populations difficult to delineate. Interestingly, all three methods assign a wide range of potency scores to RGL cells (**Figure 2F**), and the same subcluster of RGL cells is not recognized as highly potent by any method (n = 62 cells, **Figure 2E**).

### stemFinder identifies quiescent progenitors and improves annotation of scRNA-seq data

To identify the reason behind the failure of all three calculators to recognize the same RGL subcluster as highly potent, we isolate the RGL population and repeat analysis on these cells to identify two distinct subclusters. The assignment of RGL-labelled cells to subclusters corroborates stemFinder’s apparent division between more and less potent RGL subpopulations: one cluster (cluster 1) contains all but three of the cells which all methods failed to recognize as highly potent (**Figure 2G-H**).

We first attempt to determine whether cells in cluster 1 are highly potent yet quiescent. Because we previously observed the pattern of increased G1 cell proportion with increased degree of differentiation, we first examine the proportion of cells assigned to each cell cycle phase and found that cluster 1 contains a higher proportion of cells in the G1 and G2M phases than does cluster 0, suggesting that cluster 1 may be more differentiated (**Figure 2I**). We also find that both clusters exhibit a similar number of filtered transcript counts and unique features, with any low-count cells found in cluster 0, demonstrating that cluster 1 cells do not have uniquely low transcriptional activity (**Figure 2J**).

To determine gene expression differences between the RGL subclusters, we perform differential gene expression analysis and reveal that cluster 1 expresses high levels of genes also enriched in the mature astrocytes and OPCs. Genes differentially expressed in cluster 1 include S100a16 which is downregulated in multipotent stem cells differentiating into astrocytes [40-42] and the activated RGL marker Pla2g7 [43]. Cells in cluster 0 highly express genes that are enriched in nIPCs and neuroblasts, including Mycn, which plays a role in embryonic neuroblast survival and proliferation and in MEF reprogramming to iPSCs [44, 45], and Hes6, a known promoter of neuronal as opposed to astrocyte cell fate [46] (**Figure 2K**). GSEA comparing the two RGL clusters reveals that astrocyte-related terms were enriched in cluster 1 whereas cluster 0 was enriched in expression of prefrontal cortex neural progenitor markers and gene lists from progenitors from other tissue types (**Figure 2L**). Furthermore, the original UMAP plot of cell type annotations in the entire dentate gyrus dataset showed that the RGL cells which failed to be recognized as highly potent by all three methods formed a distinct cluster directly adjacent to a cluster of immature astrocytes (**Figure 2D-E**). Therefore, we propose that cluster 0 represents a true RGL progenitor whereas cluster 1 represents an activated glial progenitor primed to generate astrocyte cells.

Based on the results from stemFinder, differential expression analysis and gene set enrichment analysis (GSEA), we propose that the annotations for this dataset are revised based on our findings above (**Figure 2M**): we re-label cells from RGL cluster 1 as astrocyte progenitors with an assigned ground truth potency score of 3 (as the precursor to immature astrocytes). Under this revision, all methods continue to perform well on the entire dataset as measured by the correlation of potency score with ground truth and the discrimination accuracy of most and least differentiated populations. However, the main distinction was in percent recovery of the least differentiated cells: stemFinder was able to recover nearly all re-labeled RGL cells (95.9%) while CCAT recovered 53.1% and CytoTRACE did not recover any (0%).

We also repeat differential gene expression analysis comparing the newly defined RGL cluster to all other cells in the dataset, and detect enrichment of several neural quiescence (G0) markers in the new RGL cluster: *Scrg1* (logFC = 0.316, padjust < 0.05), *Ptprz1* (logFC = 0.747, padjust < 0.0001), *Ptn* (logFC = 1.12, *p_adjust_* < 0.0001), *Hopx* (logFC = 1.13, *p_adjust_* < 0.0001), *Sox2* (logFC = 0.480, *p_adjust_* < 0.0001), *Sox9* (logFC = 1.25, *p_adjust_* < 0.0001), *Trp53* (logFC = 0.635, *p_adjust_* < 0.0001), *Npm1* (logFC = 0.767, *p_adjust_* < 0.0001), *Mms22l* (logFC = 0.349, *p_adjust_* < 0.01), and *Tipin* (logFC = 0.912, *p_adjust_* < 0.0001). We also observed a weak yet statistically significant negative Pearson correlation between ground truth potency and neural G0 gene set score, and between stemFinder score and G0 gene set score (cor = -0.191 and cor = -0.271, *p* < 0.0001 for both comparisons). Based on these results, we hypothesize that the RGL progenitor may be in a quiescent state. This example dataset highlights the unique ability of stemFinder to identify putative quiescent progenitors, a challenge for prior methods, and to help improve upon the annotation of single-cell transcriptomic data.

### stemFinder outperforms competitors in 24 validation datasets

We validate stemFinder’s performance on 24 UMI-based single cell transcriptomic datasets spanning a variety of tissue types, developmental stages (zygote to adult), species, and sequencing platforms **(Table 1).** 16 of these datasets are from Gulati et al.; a further 17 datasets from that study were excluded as they contained lower quality data from older SC3-seq or Fluidigm C1 platforms or were derived from libraries lacking unique molecular identifiers (UMIs) which prevented our discernment of PCR duplicates from true biological signal [19, 47].

Our method performed well on the validation set as measured by the aforementioned metrics, with a mean phenotypic Spearman correlation of stemFinder score with ground truth potency of 0.561 ± 0.451, a mean single-cell Spearman correlation of 0.447 ± 0.363, a mean AUC of 0.824 ± 0.328, and a mean percent recovery of 54.7 ± 33.8%. stemFinder produces single-cell potency values with higher correlation to ground truth potency than those computed by CytoTRACE or CCAT (**Figure 3A-B**) and was significantly better at discriminating between the most and least potent populations than either competitor (**Figure 3C**). Of note, CytoTRACE also exhibits higher discrimination accuracy than CCAT in our validation set. Finally, to assess stemFinder’s ability to identify an uncharacterized population of cells with low relative degree of differentiation and high relative fate potential, we compute the percent recovery of such cells in a given dataset, as determined by a quantile-based threshold derived from the number of unique ground truth potency states in that dataset. stemFinder yielded significantly higher percent recovery of these cells (54.7 ± 33.8%) than either competitor (44.9 ± 34.9% for CytoTRACE, and 42.0 ± 31.1% for CCAT) (**Figure 3D**). Therefore, for the user who wishes to identify and characterize a progenitor population in their dataset or accurately select the root of a differentiation trajectory, stemFinder offers a reliable method for correctly identifying the least differentiated cells.

While it is helpful to compare the magnitude of performance metrics across the entire validation set, a reliable method should also be able to return the correct direction of differentiation for a single dataset, thus a binarized interpretation of performance is also appropriate. stemFinder predicted the correct direction of differentiation for 21 out of 24 (87.5%) validation datasets, as determined by a positive phenotypic Spearman correlation, where two of the failed datasets are also assigned inverted directions of differentiation by both competitor methods [48, 49]. By comparison, CytoTRACE and CCAT predicted the correct direction of differentiation for 18 out of 24 (75.0%) and 19 out of 24 (79.2%) datasets, respectively. One validation dataset, which captures the stepwise differentiation of oligodendrocyte precursors (OPCs) to mature oligodendrocytes (MOLs) in the murine CNS [50], has posed a difficulty to prior methods: CytoTRACE and CCAT return an inverted direction of differentiation for this dataset while all other methods and gene sets previously tested by Gulati et al. also performed poorly [19], yet stemFinder was able to recapitulate the correct degree of differentiation and correctly identify a fraction of OPCs as the least differentiated cells, even if its performance is not ideal (**Supplementary Figure 3A-D**).

### stemFinder is computationally efficient

To assess the computational burden of our method, we computed memory usage and run time of each potency calculator on the same validation dataset serially downsampled from *n* = 4999 filtered cells [51] (**Figure 3E-F**). Memory usage in CytoTRACE rapidly increases at *n* = 4000 cells (**Figure 3E**) while stemFinder and CCAT exhibited similar computational burden, although run times of all methods are holistically low (**Figure 3F**).

### stemFinder is robust to changes in key input parameters

The user provides two key input parameters to our method: the value of *k* to compute KNN and the cell cycle gene list. To ensure that our method is robust to alterations in these parameters, we first investigated the robustness of Gini results to changes in *k* **(Supplementary Figure 2A-D).** We computed single-cell potency for each dataset across a range of *k/k_ideal_* values, where *k_ideal_* is the rounded value of the square root of the number of cells in the filtered expression matrix. Method performance does not significantly differ by *k* ratio for any of the four performance metrics (one-way ANOVA with Tukey’s post-hoc HSD). Nonetheless, the variance in performance is minimized for 0.5 < *k/k_ideal_* < 2 and we recommend selecting a *k* value that falls within this ratio. Next, we investigated the robustness of our method to changes in cell cycle gene list. We quantify differentiation time using three different cell cycle gene lists from the following sources: Scanpy’s cell cycle gene scoring function (used in our standard pipeline), Gene Ontology, and KEGG. We found there to be no significant difference in performance, as quantified by any metric, with changes in cell cycle gene list (one-way ANOVA with Tukey’s post-hoc HSD, **Supplementary Figure 2E-H)**.

### stemFinder is robust to downsampling of the least differentiated population

To investigate the robustness of our method to downsampling of the validation dataset, we first performed random downsampling of all cells. For each validation dataset, we took a random subsample of cells from each phenotype-defined cluster at subsampling ratios 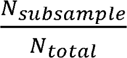 of 0.1 to 0.9. We performed proportional subsampling of each phenotype-defined cluster to ensure that the identity of the excluded cells did not disproportionately impact the performance of our method. We found no significant differences between any of the four performance metrics (one-way ANOVA and Tukey’s post-hoc HSD) for any subsampling ratios (**Supplementary Figure 2I-L)**.

We also investigate the robustness of our method to downsampling of the least differentiated cells only, or the cells with the lowest values of ground truth potency in a given dataset, which often have physiologically low abundance. Across all validation datasets, the relative abundance of these cells ranged from 0.008% to 47% with an absolute count of 1 cell to 2983 cells per dataset. We downsampled this population down to a single cell and report no significant changes observed between single-cell or phenotypic Spearman correlation or AUC with any pairwise comparison of downsampling ratios (one-way ANOVA and Tukey’s post-hoc HSD, **Supplementary Figure 2M-P).** The percent recovery of the least differentiated cells does differ when downsampled, with a paradoxically improved percent recovery seen when downsampling to the lowest ratio (0.1) compared to downsampling bins 0.7 (*p_adjust_* = 0.0451), 0.9 (*p_adjust_* = 0.0289), or 1.0 (*p_adjust_*= 0.0228). All other comparisons between downsampling bins were not significant, showing that stemFinder does not degrade in performance even if a high proportion of minimum ground truth potency cells are removed.

### Performance is optimized using cell cycle genes as opposed to randomly selected genes

As a negative control, we also tested the performance of our method given an input of randomly selected genes in lieu of our default cell cycle gene list. For each dataset, a variety of randomly selected genes were chosen with attributes (mean expression and number of cells with positive scaled expression) similar to the cell cycle genes found in that dataset. We generated *n =* 5 random gene lists per dataset and found that the performance of our method was significantly lower according to all four performance metrics if random gene lists were used as input instead of Scanpy’s cell cycle gene list **(Supplementary Figure 2M-O)**. These results further demonstrate the particular importance of cell cycle gene expression heterogeneity, rather than global gene expression heterogeneity, in computing cell potency.

### stemFinder score correlates with the proportion of cells in G1 and inversely correlates with G1, G2M, and S cell cycle gene expression

We hypothesize that predictable changes in cell cycle gene expression patterns underlie cell differentiation, either as drivers or consequences of differentiation. We first investigate the relationship between cell cycle gene expression variability, cell cycle phase assignment, and cell cycle gene set score in an example dataset in which stemFinder accurately recapitulates ground truth potency (**Figure 1C**): murine bone marrow sequenced on the 10X Genomics platform [52]. This dataset contains a cluster of Cd34+ Kit+ bone marrow hematopoeitic stem cells, or BM-HSCs (labeled by the authors as ‘stem progenitors’), as its most highly potent cell type in addition to downstream oligopotent, unipotent, and terminally differentiated cell types. GSEA of the BM-HSCs in this dataset shows enriched expression of gene lists from stem cell types across multiple tissues as compared to all other cell types in this dataset (**Figure 4A**).

To examine broad cell cycle gene expression patterns relevant to stemFinder in this dataset, we visualize the binarized, scaled expression of cell cycle genes (**Figure 4B-D**) and the cell cycle gene set score (**Figure 4E**) in single cells ordered by ascending degree of differentiation as quantified by stemFinder. Cell cycle genes exhibited both highly variable expression and high median expression in cells in the first half of this lineage, with an abrupt drop-off in cell cycle gene expression as unipotent and terminal cell types emerge. This pattern was observed across most genes from all phases of the cell cycle (S, G2M, and G1) with few exceptions, mostly in G1 (**Figure 4B-E**). Interestingly, while cell cycle gene set expression does have a negative Pearson correlation with stemFinder potency score (cor = -0.601 for S phase, cor = -0.508 for G2M phase, and cor = -0.542 for G1 phase, *p* < 0.0001 for all correlations), fitting a LOESS regression model to the data shows that cell cycle gene set scores are elevated in multipotent HSC progenitors, as expected, but they actually peak a bit later down the lineage: in the oligo-potent granulocyte, monocyte, and megakaryocyte progenitors and immature B cells. These results may support the hypothesis that quiescent cells which lowly express cell cycle genes may be better detected using cell cycle gene expression heterogeneity rather than magnitude.

Next, we examined whether high variability in cell cycle gene expression was accompanied by heterogeneity of cell cycle phase assignment in a given subpopulation. When cells in our murine bone marrow dataset were placed in order of ascending degree of differentiation, the proportion of cells in G1 increased dramatically in more differentiated cells. This pattern was present when cells are ordered by either ascending stemFinder potency score (**Figure 4F**) or by ground truth potency score (**Figure 4G**). We hypothesize that G1 is lengthened as a cell differentiates and that a longer G1 is accompanied by a decreased heterogeneity in cell cycle gene expression. These results inform us of several truths regarding cell cycle gene expression and single-cell potency: 1.) cell cycle gene expression generally increases in more potent cells yet is maximized in non-quiescent, intermediate progenitors, 2.) generalized patterns of cell cycle gene expression across a differentiation continuum are consistent across most cell cycle genes from various phases, and 3.) more differentiated subpopulations contain higher proportions of cells in the G1 phase of the cell cycle.

### Computed single-cell potency scores correlate with fate potential in a lineage tracing dataset

While many computational tools claim to compute a measure of single-cell potency, we wanted to ensure that stemFinder does not only generate scores with significant correlation to ground truth potency as defined by known lineage relationships of cell phenotypes but that undifferentiated cells with varying fate potential exhibit corresponding differences in stemFinder scores. We performed lineage tracing analysis of a hematopoietic cell differentiation dataset from Weinreb et al. in which undifferentiated cells are allowed to progress towards one of several terminal states and heritable DNA barcodes identify cells across three time points which share a common ancestor [38] (**Figure 5A-B**). Analysis of this dataset was limited only to clones containing cells in the latest time point to capture the full lineage of each undifferentiated cell. For each clonal lineage, we counted the number of distinct terminally differentiated cell types generated by an undifferentiated cell. We ran stemFinder on this data (**Figure 5C**) and found that the minimum stemFinder score of the undifferentiated cells in each clonal lineage was inversely correlated with the fate potential of that undifferentiated cell (Spearman correlation = -0.52, *p* < 0.0001, **Figure 5D**). This trend was also observed when single-cell potency was computed using CytoTRACE (Spearman correlation = -0.40, *p* < 0.0001) or CCAT (Spearman correlation = -0.47, *p* < 0.0001). We next investigated whether stemFinder score correlated with expression of marker genes of terminally differentiated cell types. We examined two cell types to this end: neutrophils (Ngp) and monocytes (Mmp8) and found a significant Pearson correlation between the stemFinder score in progenitors of these cell types or the cell types themselves and marker gene expression (cor = 0.64 and *p* < 0.0001 for neutrophil lineage and cor = 0.63, *p* < 0.0001 for monocyte lineage, respectively, **Figure 5E-F**). These findings demonstrate that 1.) stemFinder potency scores correspond to relevant biological phenomena, namely fate potential and marker gene expression, and 2.) undifferentiated cells, while annotated as one cluster, are heterogeneous in terms of cell potency and stochastic cell cycle gene expression.

## DISCUSSION

In this study, we demonstrate that neighborhood-wide variability in cell cycle gene expression is a surrogate for single-cell potency, as defined by a cell’s degree of differentiation and fate potential. We contrast this definition with other related concepts such as developmental stage, imminent propensity for cell division, and self-renewal capacity, for our method recognizes both adult stem cells and quiescent cells as highly potent. While we demonstrate stemFinder’s ability to distinguish subpopulations of undifferentiated cells which generate different numbers of terminally differentiated cell types in a lineage tracing dataset [38], most validation datasets are not designed to analyze fate potential. Nonetheless, stemFinder performs well in other validation datasets where ground truth potency is dictated by stepwise progression from least to most differentiated cell types in one or more lineages.

The methodology presented here, available in the R package stemFinder, is robust, exhibits equal or superior performance to prior methods as measured by several metrics in a cohort of UMI-based scRNA-seq data, and theoretically would be applicable to any species for which paralogs of the cell cycle gene. stemFinder may be used in lieu of marker genes and a priori cell type information to identify rare and putatively highly potent progenitors or in lieu of pseudotime to align cells along a differentiation trajectory. Our method can also be used to study cell cycle gene expression dynamics across cell differentiation in multiple contexts to investigate the underlying biological role of the cell cycle in differentiation.

We extend the applicability of stemFinder in two ways beyond that of the traditional potency calculation method: first, in the refinement of existing cell type annotations and clustering in scRNA-seq data in complement with traditional gene expression analysis techniques, and second, in the investigation of patterns of cell cycle phase and cell cycle gene expression. We also demonstrate stemFinder’s utility in revealing biologically relevant heterogeneity in cell types thought to be transcriptionally homogeneous, as in the murine dentate gyrus RGL population [39].

In future work, we suggest the application of stemFinder to a validation dataset designed to further dissect out its relationships to fate potential and differentiation time. To validate our findings that cell cycle gene expression heterogeneity inversely correlates with the proportion of cells in the G1 phase of the cell cycle, possibly due to G1 lengthening with differentiation [32], we suggest in vitro studies where time lapse imaging of cell cycle gene expression is analyzed alongside barcoded single-cell transcriptomic data. Finally, our benchmarking depends on the numeric approximation and definition of “ground truth,” and we hope for this definition to become even clearer as datasets are generated with this biological question in mind.

## CODE AVAILABILITY

stemFinder is available as an R package at https://github.com/cahanlab/stemfinder and scripts to reproduce all analyses are available at https://github.com/CahanLab/stemfinder_paper.

## DATA AVAILABILITY

No new data were generated from this study. All data used in this study are publicly available as shown in **Table 1**.

---

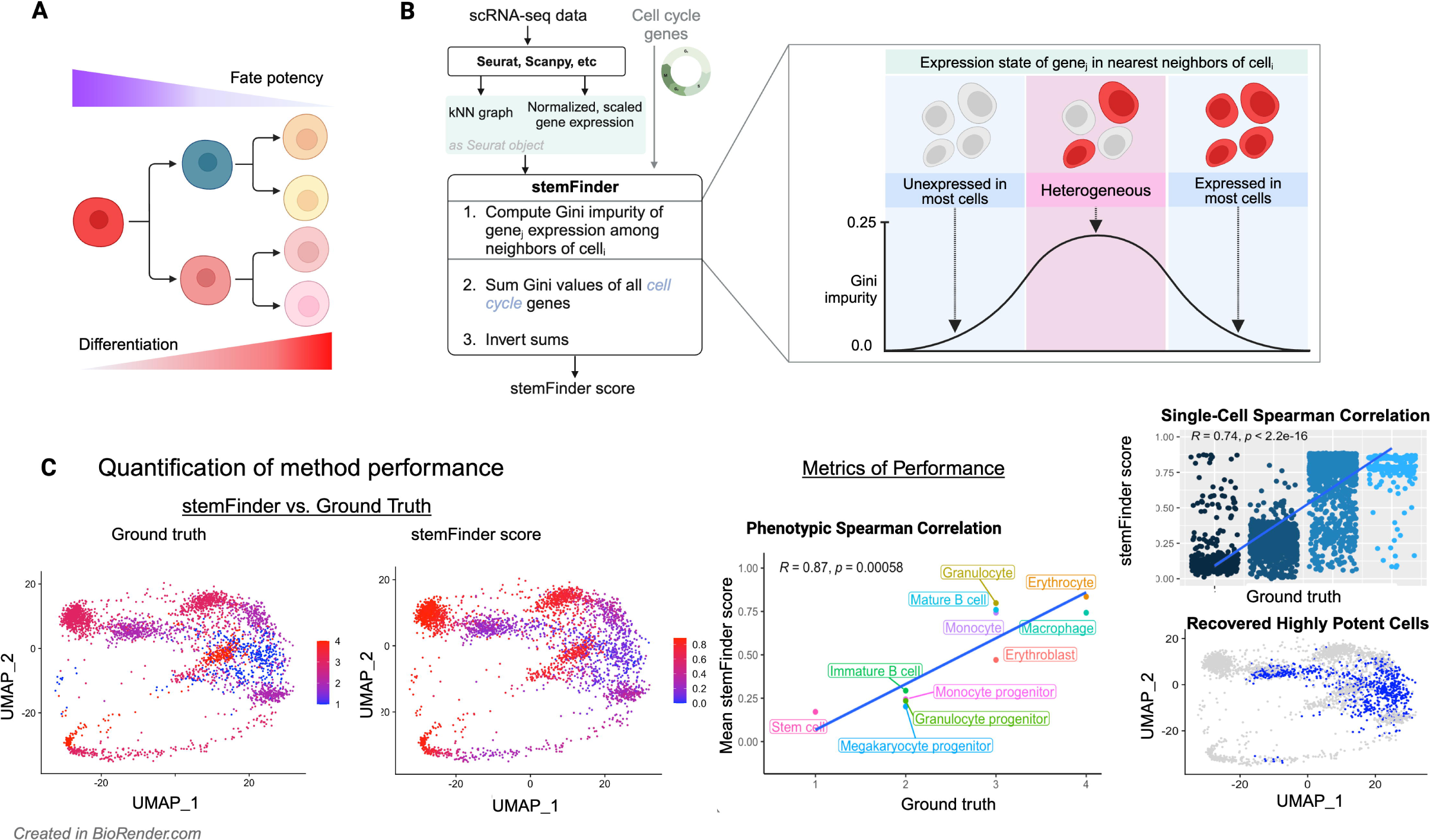

| Validation data | Platform | Species | Tissue type | GEO Accession | From Gulati et al. validation or training set? | Timepoints? | Number of filtered cells | Number of cells with highest ranked potency | Percentage known highly potent cells |
| --- | --- | --- | --- | --- | --- | --- | --- | --- | --- |
| Mouse 10X bone marrow | 10X | *Mus musculus* | Bone marrow, adult | GSE109774 | Yes | No | 3427 | 399 | 11.6428363 |
| *C elegans* muscle mesoderm 10X | 10X | *C. elegans* | Mesoderm, embryonic | GSE126954 | Yes | No | 22370 | 21 | 0.093875726 |
| Dentate gyrus 10X mouse phenotype | 10X | *Mus musculus* | Brain, adult | GSE95753 | Yes | No | 5344 | 176 | 3.293413174 |
| Endometrium CEL-seq mouse | CEL-seq | *Mus musculus* | Uterus, adult | GSE98451 | Yes | Yes | 8538 | 173 | 2.026235652 |
| *C elegans* hypoderm 10X | 10X | *C. elegans* | Hypodermis, embryonic | GSE126954 | Yes | No | 12249 | 1 | 0.008163932 |
| Direct programming in vitro neuron inDrop | inDrop | *Mus musculus* | Spinal motor neuron and ESC, *in vitro* | GSE97391 | Yes | No | 2366 | 1109 | 46.87235841 |
| Standard protocol in vitro neuron inDrop | inDrop | *Mus musculus* | Spinal motor neuron and ESC, *in vitro* | GSE97391 | Yes | No | 2411 | 956 | 39.65159685 |
| Lgr5+ intestine CEL-seq mouse | CEL-seq | *Mus musculus* | Small intestine, adult | GSE76408 | Yes | No | 480 | 131 | 27.29166667 |
| Peripheral blood 10X human | 10X | *Homo sapiens* | Blood, adult | SRP073767 | Yes | No | 4999 | 500 | 10.0020004 |
| Whole intestine Drop-seq mouse Haber et al. | Drop-seq | *Mus musculus* | Small intestine, adult | GSE92332 | Yes | No | 4581 | 830 | 18.11831478 |
| Dentate gyrus 10X mouse timepoints | 10X | *Mus musculus* | Brain, adult | GSE95753 | Yes | Yes | 6000 | 578 | 9.633333333 |
| Neural stem cells, Drop-seq mouse | Drop-seq | *Mus musculus* | Brain, embryonic | GSE107122 | Yes | Yes | 5925 | 1999 | 33.73839662 |
| Thymus Drop-seq mouse | Drop-seq | *Mus musculus* | Thymus, embryonic | GSE107910 | Yes | Yes | 9307 | 361 | 3.878800903 |
| Gdf5+ joint stem cells, day 0-4, 10X mouse | 10X | *Mus musculus* | Synovial joint, embryonic | GSE151985 | No | Yes | 7329 | 2632 | 35.91212989 |
| Hair epidermis, C1 mouse | C1 | *Mus musculus* | Epidermis, adult | GSE67602 | Yes | No | 720 | 330 | 45.83333333 |
| Oligodendrocyte precursor phenotypes, C1 mouse | C1 | *Mus musculus* | CNS, adult | GSE75330 | Yes | No | 4974 | 308 | 6.192199437 |
| Pancreatic acinar cells 10X mouse | 10X | *Mus musculus* | Pancreas, adult | GSE172380 | No | No | 6079 | 546 | 8.981740418 |
| hiPSC differentiation to cardiomyocyte, SPLiT-seq human | SPLiT-Seq | *Homo sapiens* | iPSC to cardiomyocyte, *in vitro* | On Quiltdata: see PMID 34349150 | No | No | 18523 | 814 | 4.394536522 |
| mESC directed differentiation, MULTI-seq | MULTI-seq | *Mus musculus* | ESC to mesoderm / neurectoderm / endoderm, *in vitro* | GSE177051 | No | Yes - but ground tru | 5228 | 1071 | 20.48584545 |
| Heurault et al. Aging HSC, 10X mouse | 10X | *Mus musculus* | Bone marrow, adult (2mo old mice only) | GSE147729 | No | No | 3794 | 642 | 16.92145493 |
| Skeletal muscle progenitors, 10X human | 10X | *Homo sapiens* | Skeletal muscle, adult | GSE143704 | No | No | 4671 | 2983 | 63.86212802 |
| Oviduct, mouse P4 and adult, 10X | 10X | *Mus musculus* | Oviduct, adult | GSE180102 | No | Yes | 1070 | 263 | 24.57943925 |
| Zebrafish embryo timepoints, Drop-seq | Drop-seq | *Danio rerio* | Embryo | GSE106587 | Yes | Yes | 6000 | 54 | 0.9 |
| NAFLD hepatocytes and non-parenchymal cells, 10X mouse | 10X | *Mus musculus* | Liver, adult | GSE166504 | No | No | 8966 | 409 | 4.561677448 |
| Test data | Platform | Species | Tissue type | GEO Accession | Notes |  |  |  |  |
| Lineage tracing hematopoiesis | inDrop | *Mus musculus* | HPCs, *in vitro* | GSE140802 | Cells transduced with lineage barcodes prior to culture |  |  |  |  |

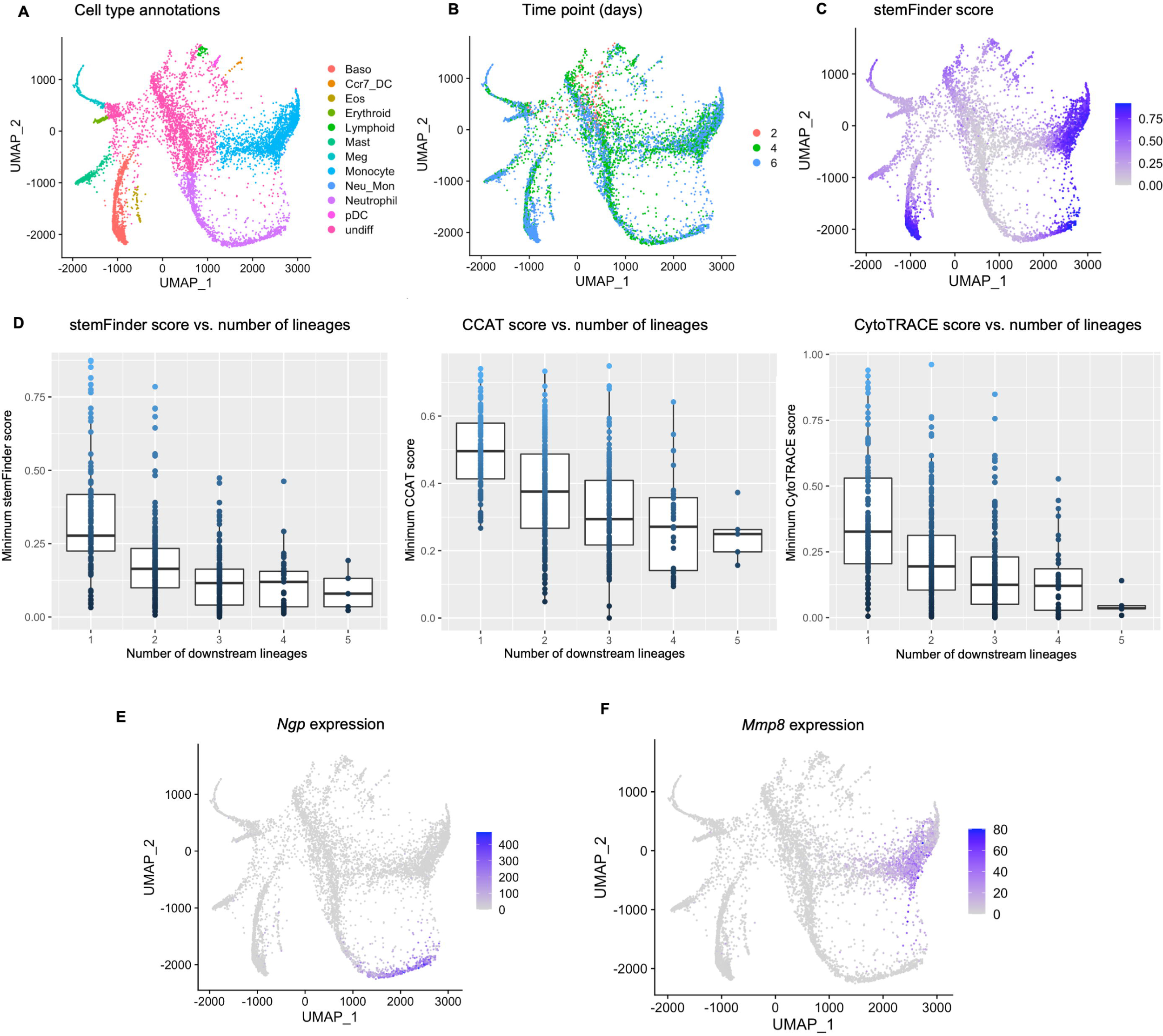

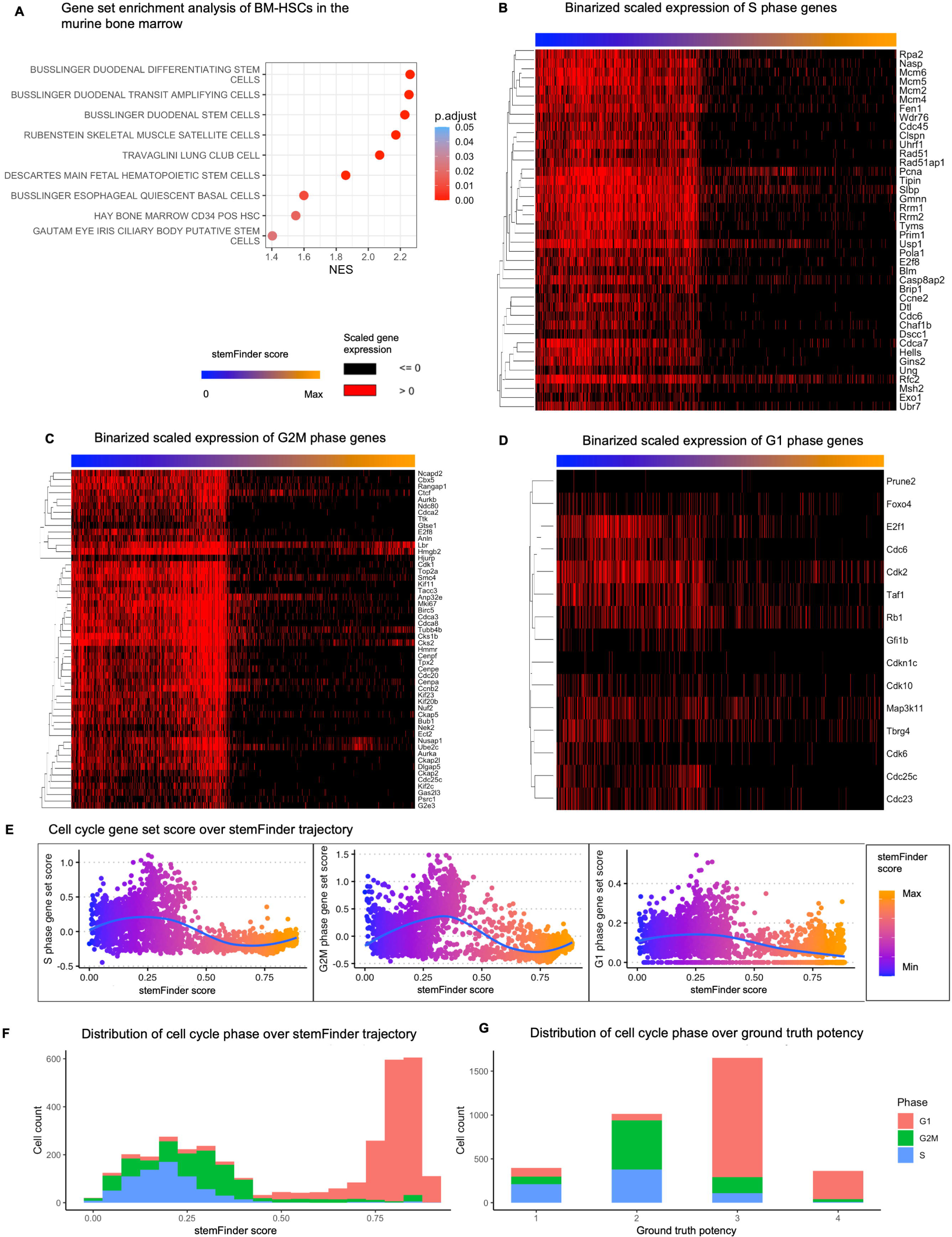

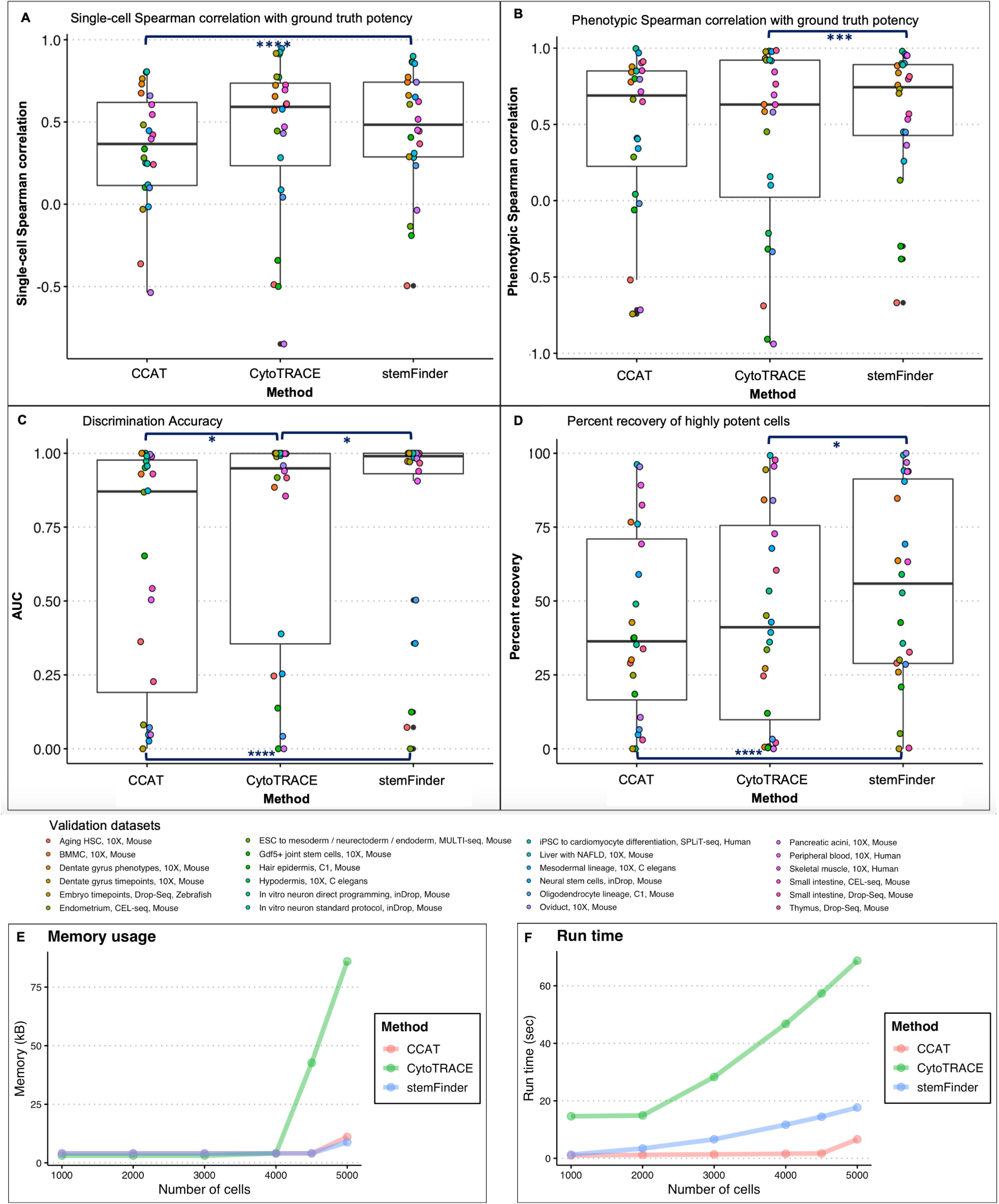

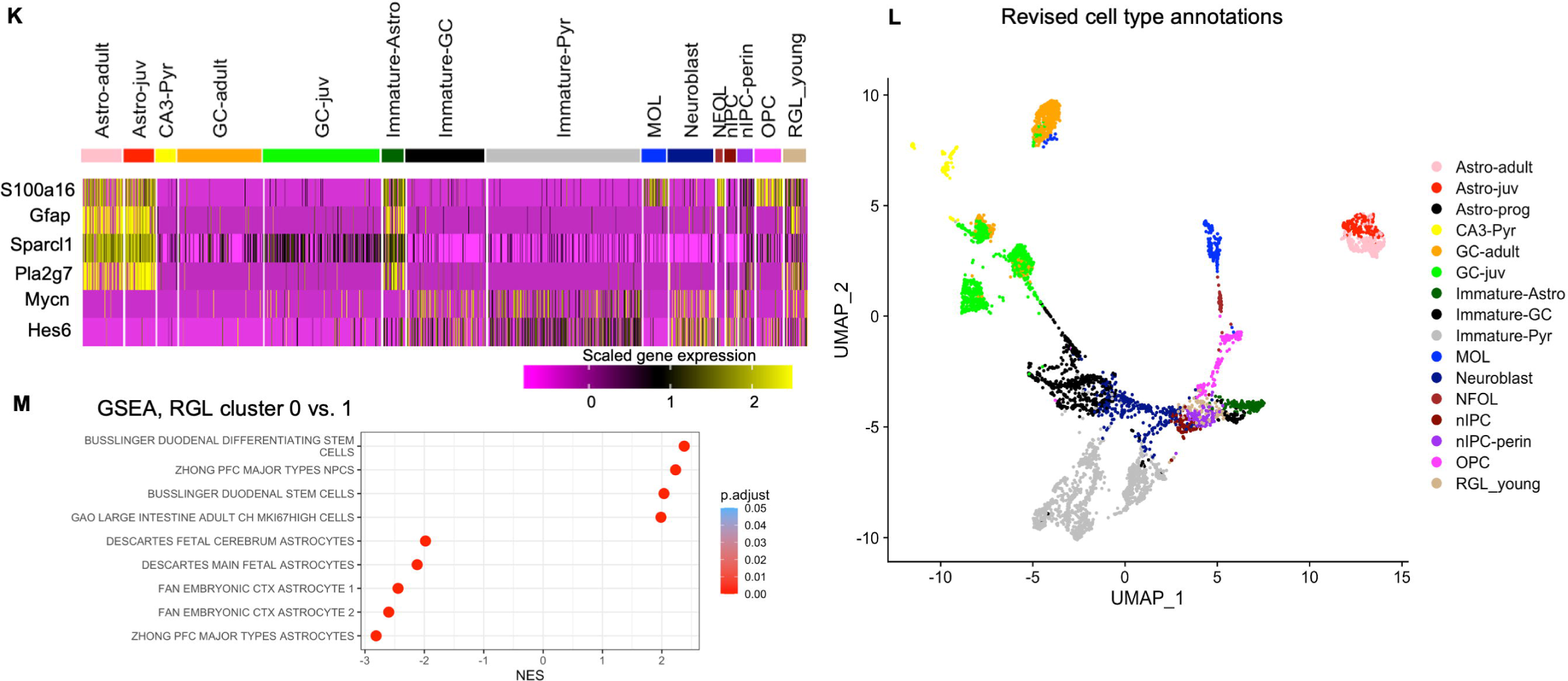

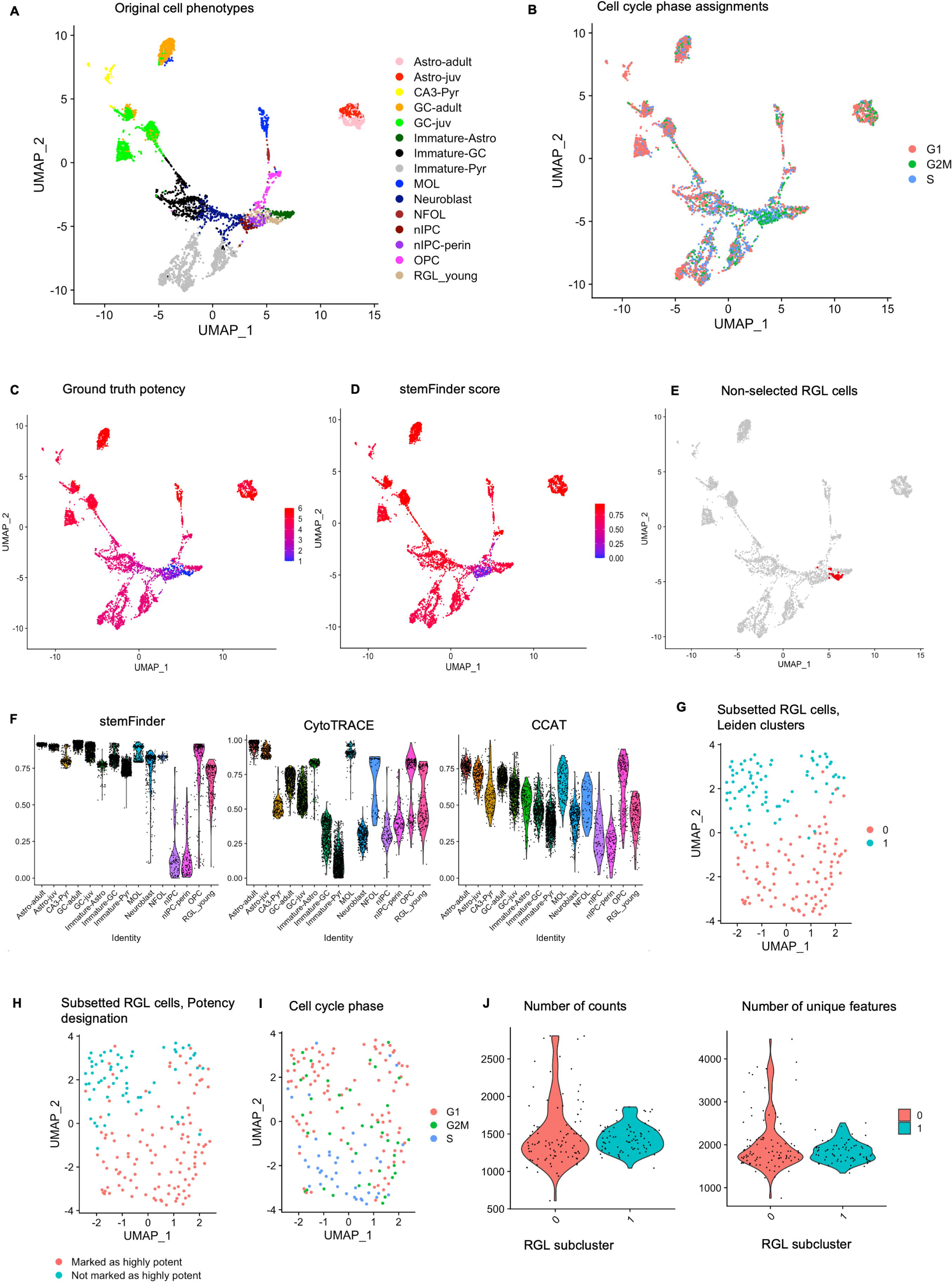

