## Supplementary material for "Cell cycle expression heterogeneity predicts degree of differentiation": Legends

**MAIN FIGURE LEGENDS**

**Figure 1** Overview of stemFinder

**(A)** The goal of stemFinder is calculate single cell estimates of the extent of differentiation or fate potency from single cell RNA-seq (scRNA-seq) data.

**(B)** stemFinder takes as input a Seurat object containing log-normalized, scaled gene expression estimates, a K-nearest neighbors graph which is straightforward to generate in R or Python, and a list of cell cycle genes which are provided for human, mouse, and *C. elegans*. stemFinder computes a metric of variability in cell cycle gene expression for each gene in each query using Gini impurity. To derive differentiation time estimates, Gini impurity values are summed across cell cycle genes within each cell and then inverted such that a value of zero corresponds to a cell with high fate potential or low degree of differentiation, and a maximum possible score of 1 corresponds to a cell with relatively low fate potential or high degree of differentiation in that dataset.

**(C)** The performance of differentiation time predictors such as stemFinder are assessed with four metrics, three of which are illustrated here, in comparison to ground truth fate potency or developmental time as defined by known lineage relationships of cell types or developmental time points, depending on the validation dataset.

**Figure 2** Application of stemFinder for improved annotation of the murine dentate gyrus.

**(A-B)** Original cell phenotype annotations (A) and cell cycle phase assignments (B) of murine dentate gyrus cells on UMAP embedding.

**(C-D)** Single-cell values of ground truth potency, represented as differentiation time based on known lineage relationships of each cell phenotype (C), and stemFinder score (D) on UMAP embedding in the murine dentate gyrus.

**(E)** UMAP embedding of all cells in the murine dentate gyrus validation dataset, where cells highlighted in red are RGL cells, or the cell phenotype with the lowest ground truth differentiation time, which were not selected as highly potent by any of the three potency calculators (stemFinder, CytoTRACE, or CCAT).

**(F)** Violin plots of single-cell scores computed by stemFinder, CytoTRACE, and CCAT grouped by cell phenotype.

**(G-I)** Subsetted and re-clustered RGL cells, the least differentiated cell type in the murine dentate gyrus dataset, on UMAP embedding. Cells are colored by Leiden cluster assignment (G), by whether they are marked as highly potent by any potency calculator (H), and by cell cycle phase assignment (I).

**(J)** Violin plot depicting the number of filtered transcript counts (left) and filtered features (right) per cell in RGL cells, grouped by RGL subcluster.

**(K)** Heat map of gene expression levels of select differentially expressed genes between RGL cluster 0 versus RGL cluster 1. Gene expression is shown in all cells from the entire murine dentate gyrus validation dataset.

**(L)** Dot plot of select gene set enrichment analysis results comparing genes with upregulated expression in RGL cluster 0 versus RGL cluster 1. Normalized enrichment score (NES) > 0 indicates a term which is enriched in the evaluated subpopulation and vice versa. All terms have adjusted p value < 0.05, where dots are colored by adjusted p value.

**(M)** Revised cell type annotations on UMAP embedding for murine dentate gyrus cells. A new cell type, astrocyte progenitors (“Astro-prog”) was created based on gene set enrichment analysis, subclustering, and stemFinder results.

**Figure 3** stemFinder outperforms prior potency quantification methods for UMI-based scRNA-seq data.

**(A)** Spearman correlation between single-cell values of ground-truth potency and potency quantified by one of three methods: CCAT, CytoTRACE, and our method (stemFinder). Data points are colored by dataset, which is labelled by cell type, sequencing platform, and organism (see legend). **(B)** Spearman correlation between mean ground-truth potency for each phenotype-defined cluster and mean quantified potency for each phenotype-defined cluster across all methods. **(C)** Discrimination accuracy (AUC), or the ability of a method to distinguish between the potency of the most and least potent populations, across all methods. **(D)** Percent recovery of highly potent cells by each method. **(E-F)** Memory usage in kB (E) and total elapsed run time in seconds (F) for our method as compared to prior methods, as tested on the peripheral blood 10X human query dataset, which contains *n =* 4999 filtered cells.

**** *p* < 0.0001, *** *p* < 0.001, ** *p* < 0.01, * *p* < 0.05

**Figure 4** Characteristic cell cycle gene expression patterns underlie cell differentiation.

**(A)** Gene set enrichment analysis using the MSigDB C8 reference database for annotated BM-HSCs in the murine bone marrow 10X validation dataset. Enriched terms have NES > 0 and adjusted p value < 0.05.

**(B-D)** Heat map of binarized scaled gene expression in the murine bone marrow: S phase genes (B), G2M phase genes (C) and G1 phase genes (D). Cells are arranged on the X axis in order of ascending stemFinder score. The dendrogram on the y axis depicts results of hierarchical clustering of genes by Pearson correlation of gene expression values.

**(E)** Scatter plot of single-cell values of stemFinder score and gene set score (for S, G2M, or G1 phase genes) in the murine bone marrow with a fitted LOESS regression model.

**(F-G)** Stacked bar plots depict the number of cells with a given binned potency value, either stemFinder score (F) or ground truth potency (G), belonging to the G1, G2M, or S phases of the cell cycle.

**Figure 5** stemFinder scores correlate with fate potential in a lineage tracing dataset.

**(A-C)** Cell type annotations (A), time points (B), and stemFinder scores (C) for a hematopoietic cell lineage tracing dataset on UMAP embedding.

**(D)** Box plot of the number of downstream terminally differentiated cell types in a single clonal lineage versus the minimum stemFinder score, CCAT score, and CytoTRACE score of undifferentiated cells within a single clonal lineage.

**(E-F)** Gene expression of neutrophil (E) and monocyte (F) marker genes on UMAP embedding.

**SUPPLEMENTARY FIGURE LEGENDS**

**Supplementary Figure 1** Heterogeneity of cell cycle gene expression inversely correlates with ground truth potency.

Single-cell Spearman correlation of ground truth potency and stemFinder potency score, a metric of cell cycle gene expression heterogeneity, or inverted cell cycle gene set score, respectively. Each point corresponds to one validation dataset (*n* = 24) and points are colored by input gene list type: G2M + S phase genes (standard input to stemFinder), G1 + G2M + S phase genes, G1 phase only, G2M phase only, and S phase only genes **(A).** Four performance metrics compare the performance of stemFinder, which computes the heterogeneity of cell cycle (G2M and S phase) gene expression, to the gene set score for G2M and S phase genes in each validation dataset **(B)**. Note that the gene set score is inverted so that a positive correlation with ground truth represents a correct reconstruction of ground truth potency.

**** *p* < 0.0001, *** *p* < 0.001, ** *p* < 0.01, * *p* < 0.05

**Supplementary Figure** 2 stemFinder is robust to changes in *k*, cell cycle gene lists, and downsampling.

**(A-D)** Results of robustness testing for *k* as input to K-nearest neighbors. We measure robustness as deviation from performance at *k*_ideal_, or the square root of the number of cells in the filtered expression matrix. *k* values are expressed as ratios of *k*/*k*_ideal_, with each box corresponding to one *k*/*k*_ideal_ bin. Performance metrics measured are single cell Spearman correlation (A) and phenotypic Spearman correlation (B) with ground truth potency, AUC (C), and percentage recovery of highly potent cells (D). Results are reported as the difference in performance between a given value of the x-axis variable and the default value of the x-axis variable (here, *k*/*k*_ideal_ = 1).

**(E-H)** Results of robustness testing for different input cell cycle gene lists to stemFinder: Regev lab cell cycle gene list (default input, provided in stemFinder), KEGG cell cycle gene list, and GO cell cycle gene list. Robustness is measured as deviation from performance given the standard Regev lab input gene list in terms of single-cell Spearman correlation (E) and phenotypic Spearman correlation (F) with ground truth potency, AUC (G), and percentage recovery of highly potent cells (H).

**(I-L)** Results of robustness testing for random downsampling of the entire population, where results are binned by ratio of the original number of cells retained in the downsampled dataset. We measure robustness as deviation from performance when all cells in the original filtered dataset are retained for analysis. Performance metrics measured are single cell Spearman correlation (I) and phenotypic Spearman correlation (J) with ground truth potency, AUC (K), and percentage recovery of highly potent cells (L).

**(M-P).** Results of robustness testing for downsampling of the most potent ground truth population, where results are binned by percentage of the original potent population retained in the downsampled dataset. We measure robustness as deviation from performance when all cells in the original filtered dataset are retained for analysis. Performance metrics measured are single cell Spearman correlation (M) and phenotypic Spearman correlation (N) with ground truth potency, AUC (O), and percentage recovery of highly potent cells (P).

**(Q-T).** Comparison of performance of our method with two different input gene lists: Regev lab cell cycle genes versus an equal number of randomly selected genes with similar expression profiles. Performance is quantified as single-cell Spearman correlation (Q) and phenotypic Spearman correlation (R) with ground truth potency, AUC (S), and percentage recovery of highly potent cells (T).

**** *p* < 0.0001, *** *p* < 0.001, ** *p* < 0.01, * *p* < 0.05

**Supplementary Figure 3.** stemFinder returns the correct direction of differentiation in the murine oligodendrocyte lineage.

**(A)** Cell type annotations on UMAP embedding for the murine oligodendrocyte C1 validation dataset. **(B)** Violin plot of stemFinder score grouped by cell phenotype in the oligodendrocyte dataset. **(C)** Violin plot of CytoTRACE (left) and CCAT (right) potency scores grouped by cell phenotype in the oligodendrocyte dataset. **(D)** Ground truth potency, as represented by differentiation time based on known lineage relationships of cell phenotypes, and scores from stemFinder, CytoTRACE, and CCAT on UMAP embedding.
