## Supplementary Methods for "Cell cycle expression heterogeneity predicts degree of differentiation"

***scRNA-seq data processing***

To obtain a filtered, log-normalized, and scaled Seurat object for each validation dataset, we first obtain the raw counts data provided on GEO and create a Seurat object from the counts matrix and any provided metadata (version 5.1.0) [1]. Each cell in the Seurat object has a minimum number of 200 expressed features and each feature is expressed in a minimum number of 5 cells. We compute percent mitochondrial and ribosomal gene expression and remove mitochondrial genes and *Malat1* from the dataset. We filter out putative doublets by subsetting for cells with number of features less than *f_max_*, or the 95^th^ quantile of nFeature_RNA for the dataset. We further remove cells with percent mitochondrial gene expression less than 15% by default, though a violin plot of percent mitochondrial gene expression was examined for each dataset to perform adaptive thresholding and remove outliers. Next, we perform log normalization with the NormalizeData() function in Seurat, compute highly variable genes with FindVariableFeatures() (variance stabilizing transformation method, *n =* 2500 features), and scale all features in the resulting matrix with the ScaleData() function. We remove any variable features that are also contained in the cell cycle gene list so that cell cycle gene expression does not influence downstream neighbor assignments. We also perform cell cycle scoring using S genes and G2M genes from the Scanpy cell cycle scoring gene list. Finally, the FindNeighbors() function in Seurat is used to construct a K-nearest neighbors (KNN) graph based on the Euclidean distance in PCA space. The value of *k* is chosen as the square root of N, or the number of cells in the dataset, which is a standard practice in scRNA-seq data analysis [2]. The number of principal components was chosen from visual inspection of the elbow plot. KNN is a commonly used machine-learning method which draws edges between cells to form a graph, where cells with similar gene expression patterns are connected [3]. This step returns a *cell x cell* matrix where the connection between cell pairs is represented by a binary entry.

***Cell phenotype annotation and selection of ground truth potency***

If metadata were not provided by the original authors, Leiden clustering was performed in Seurat and clusters were annotated as directed by the original study. Cell phenotype annotations were validated using known marker gene expression and differential gene expression analysis (Wilcox rank sum test, FindAllMarkers() in Seurat).

Numeric ground truth potency values were assigned to single cells, where the lowest ground truth potency value corresponds to the least differentiated cell(s) in the dataset and the highest ground truth potency value corresponds to the most differentiated cell(s) in the dataset. For time series datasets (*n = 12* validation datasets), time points of sample acquisition were used as ground truth potency values unless otherwise noted in **Table 1**. For all other datasets, ground truth potency values were assigned based on cell phenotype and established lineage relationships.

For datasets provided on the CytoTRACE website (<https://cytotrace.stanford.edu/>), our data processing pipeline is the same as above except where noted below and is detailed in our process_cyto_adata() function. UMAP coordinates and metadata are provided by the authors for each dataset; metadata contains cell phenotype and ground truth potency values. Cells are not filtered by maximum number of features or percent mitochondrial gene expression as above; instead, cells with annotated phenotypes are selected as they have already passed quality control. Ground truth potency values assigned by Gulati et al. were upheld for fair comparison [4].

***Analysis of lineage tracing LT-scSeq dataset***

The gene expression matrix, cell type annotations, time points, embeddings, and clonal labels were provided by the original authors [5]. The X_clone matrix was used to identify members of a given clonal lineage. To determine the number of terminally differentiated cell types in each clonal lineage, or its fate potential, the number of unique cell types (not including “undifferentiated” cells) per clonal lineage were counted. For each clonal lineage, the minimum stemFinder score of the undifferentiated cells within that lineage was recorded. Pearson correlations between minimum stemFinder score and fate potential were determined using the cor.test() function in the base *stats* package in R.

***Potency quantification with CytoTRACE and CCAT***

The CytoTRACE R package (version 0.3.3) was installed using devtools (http://cytotrace.stanford.edu). CytoTRACE results are computed on raw, unfiltered counts matrices in R using the CytoTRACE() function without subsampling. Resulting CytoTRACE values range from 0 (more differentiated) to 1 (more potent).

The SCENT R package, which contains the functions for CCAT, was installed using devtools (https://github.com/aet21/SCENT). CCAT potency values were computed on normalized and log-transformed counts matrices. Feature names were converted to ENTREZ IDs using gprofiler2 (version 0.2.1, CRAN) prior to running CCAT with the CompCCAT() function in R. CCAT returns a potency score ranging from 0 (more differentiated) to 1 (more potent).

Inverted CytoTRACE and CCAT scores were computed as $[1 - score/max(score)]$ to correspond to ground truth potency and to stemFinder directionality, where lower values correspond to less differentiated cells.

***Differentiation quantification with stemFinder***

The code for our method is written in R (“run_stemFinder”) and takes the following inputs: 1.) a Seurat object containing log normalized and scaled scRNA-seq data, 2.) a nearest-neighbors matrix with dimensions of *cell x cell*, 3.) a numeric value for k, the parameter used as input to compute the nearest-neighbors matrix, 4.) a numeric threshold for binning the scaled gene expression matrix (default: zero), and 5.) a list of marker genes expressed in the filtered query data. We provide a reference list of S and G2M cell cycle genes for 3 species: *Homo sapiens, Mus musculus,* and *C. elegans*. The cell cycle gene list is derived from the Scanpy package for scRNA-seq analysis in Python [6] and orthologs were obtained using the gprofiler2 R package [7].

Our method performs the following steps to compute differentiation time provided the inputs above:

1. Subset the scaled gene expression matrix to contain only the gene expression information for inputted marker genes (default: S and G2M phase cell cycle genes). Binarize scaled expression values for each gene in the subsetted matrix using the inputted threshold (default: zero scaled expression).
2. Compute Gini impurity for each gene in each cell. The Gini impurity will serve as a measure of stochasticity of the expression of that gene in the query cell as compared to the (*k* – 1) non-query cells in its surrounding neighborhood (the cells with similar transcriptomic profiles to the cell of interest; presumably the same cell type; note that Seurat marks the query cell as one of its own *k* nearest neighbors). Gini impurity will be zero if the expression of a given gene in a given neighborhood is homogenous (either with scaled expression above the given threshold for all non-query cells in a given neighborhood, or with scaled expression at or below the given threshold for all non-query cells in a given neighborhood) and it will be maximized at 0.25 if the expression of that gene in that neighborhood is heterogeneous (50% probability of the gene having a scaled expression above the given threshold in a neighboring cell).
   a. Gini impurity for each query cell $G_{i}$ is computed according to the following formula:
      i. $G_{i}=\sum_{j} p_{ij}\times(1-p_{ij})$
      ii. Where $p_{ij}=\frac{n}{(k-1)}$ is the probability that the binarized expression pattern of gene *j* in a neighborhood matches that of the query cell *i*
      iii. Where $n$ is the number of non-self neighboring cells with the same expression pattern of a given gene as the cell of interest, $k$ is the number of neighbors used to compute KNN, $i$ is the query cell, and $j$ is the given gene
   b. The Gini values across all genes are summed for each cell to obtain a per-cell raw stemFinder score. The resulting Gini value is then divided by the maximum Gini value for that dataset and subtracted from one to obtain an inverted score which ranges from 0 (least differentiated) to a maximum possible score of 1 (most differentiated).
   c. Both stemFinder scores are stored as metadata columns (“stemFinder_raw” and “stemFinder”) in the returned Seurat object.

***Evaluation of differentiation time predictors***

We quantify the performance of each potency calculator across *n =* 5 iterations per validation dataset in terms of phenotypic Spearman correlation, single-cell Spearman correlation, AUC, and percent recovery of potent cells, as outlined in the functions compute_performance_single(), auc_probability(), and pct_recover() on stemFinder GitHub. Phenotypic Spearman correlation is defined as the Spearman correlation between the mean ground truth potency of each phenotype-defined cluster and the mean computed potency of each phenotype-defined cluster in a given dataset. Single-cell Spearman correlation is defined as the Spearman correlation between individual cellular values of ground truth potency and individual cellular computed potency values for all cells in a given dataset. AUC quantifies the ability of a potency calculator to discriminate between the most and least potent ground truth cells in a given dataset. The compute_performance_single() function takes as input a Seurat object containing metadata columns of scores from stemFinder and optionally from a competitor method, a logical indicating if a comparative analysis between competitor method and ground truth is desired, and a logical indicating if competitor scores are inverted to correspond to ground truth, stemFinder, and pseudotime directionality. It returns a list containing the single-cell and phenotypic Spearman correlations and AUC for stemFinder and optionally a competitor method for that dataset.

Percent recovery is computed as the percentage of cells in the lowest ground truth potency cluster which are given a stemFinder score below a quantile-based threshold. The quantile-based threshold is assigned to each dataset as $\frac{1}{N_{GT}}$, where N_GT_ is the number of unique ground truth potency states in that dataset. Percent recovery can be computed using the function pct_recover() which takes as input a Seurat object containing ground truth potency values and stemFinder scores for a query scRNA-seq dataset and returns a numeric value of percent recovery.

***Robustness testing***

We perform robustness testing for our method across all validation datasets to quantify changes in performance, as measured by single-cell and phenotypic Spearman correlations, AUC, and percent recovery of highly potent cells with changes in key parameters: number of nearest neighbors *k,* downsampling of the highly potent population, downsampling of cells across all phenotypes, and changes in input marker gene list.

To test robustness to changes in *k,* we test a range of *k/k_ideal_* ratios from 0.2 to 4.0 with a minimum of *k =* 2, where *k_ideal_* is the rounded square root of the number of cells in the filtered dataset. stemFinder is run for *n =* 3 iterations per *k* value per dataset. To test robustness to downsampling of the highly potent population, we randomly sample (without replacement) cells from the population containing the lowest ground truth potency score at fixed ratios ranging from 0.1 to 0.9 in increments of 0.1. We run stemFinder on the downsampled dataset containing all cells with ground truth potency > min(ground truth potency) and the downsampled highly potent cells for *n =* 3 iterations per downsampled dataset. To test robustness to downsampling of cells across all phenotypes, we randomly sample (without replacement) cells from each phenotype-defined cluster at fixed ratios ranging from 0.1 to 0.9 in increments of 0.1 and run stemFinder for *n =* 3 iterations per downsampled dataset. To test robustness to changes in input marker gene list, we first compare performance across 3 cell cycle gene lists: Regev lab G2M and S genes (default), Gene Ontology cell cycle genes, and KEGG cell cycle genes for a series of *n =* 3 iterations per gene list per dataset. We also compare performance of Regev cell cycle genes to randomly selected genes, where random sampling (without replacement) of genes from the filtered dataset was performed to generate *n =* 5 random gene lists of equal length to the cell cycle genes per dataset.

For all robustness testing, the stemFinder method and performance quantification are performed as detailed above. We compute the negative deviation in performance for each *k* value, downsampling ratio, or alternative gene list as compared to the performance at default values of *k =* $\sqrt{N_{cells}}$, downsampling ratio = 1.0, or gene list = Regev lab G2M and S cell cycle genes, respectively.

To perform an assessment of run time and memory usage, each method was run on the same query dataset on a Ubuntu c5.4xlarge S3 instance. The query dataset was randomly downsampled from *n =* 4999 total cells to *n =* 1000, 2000, 3000, 4000, and 4500 cells.

***Statistical analysis***

To compute Spearman correlations between ground truth and computed potency, we use the cor.test() function from the base *stats* package in R (version 4.0.3). To compute discrimination accuracy, or AUC, we use the auc_probability() function in R [8]. To compare performance metrics between different methods, we employ the pairwise T test with Bonferroni correction and use the significance threshold of adjusted p value < 0.05 (pairwise_t_test() function from the *rstatix* package in R, version 0.7.0). To compare results of robustness testing, we use one-way ANOVA and Tukey’s post-hoc HSD with a significance threshold of adjusted p value < 0.05 (aov() and TukeyHSD() functions from the base *stats* package in R). To compare the performance of stemFinder using cell cycle versus random gene lists, we use the unpaired T test (t.test() from the base *stats* package in R).
