## Supplementary figures and images for "Cell cycle expression heterogeneity predicts degree of differentiation"

### FigS1.tiff

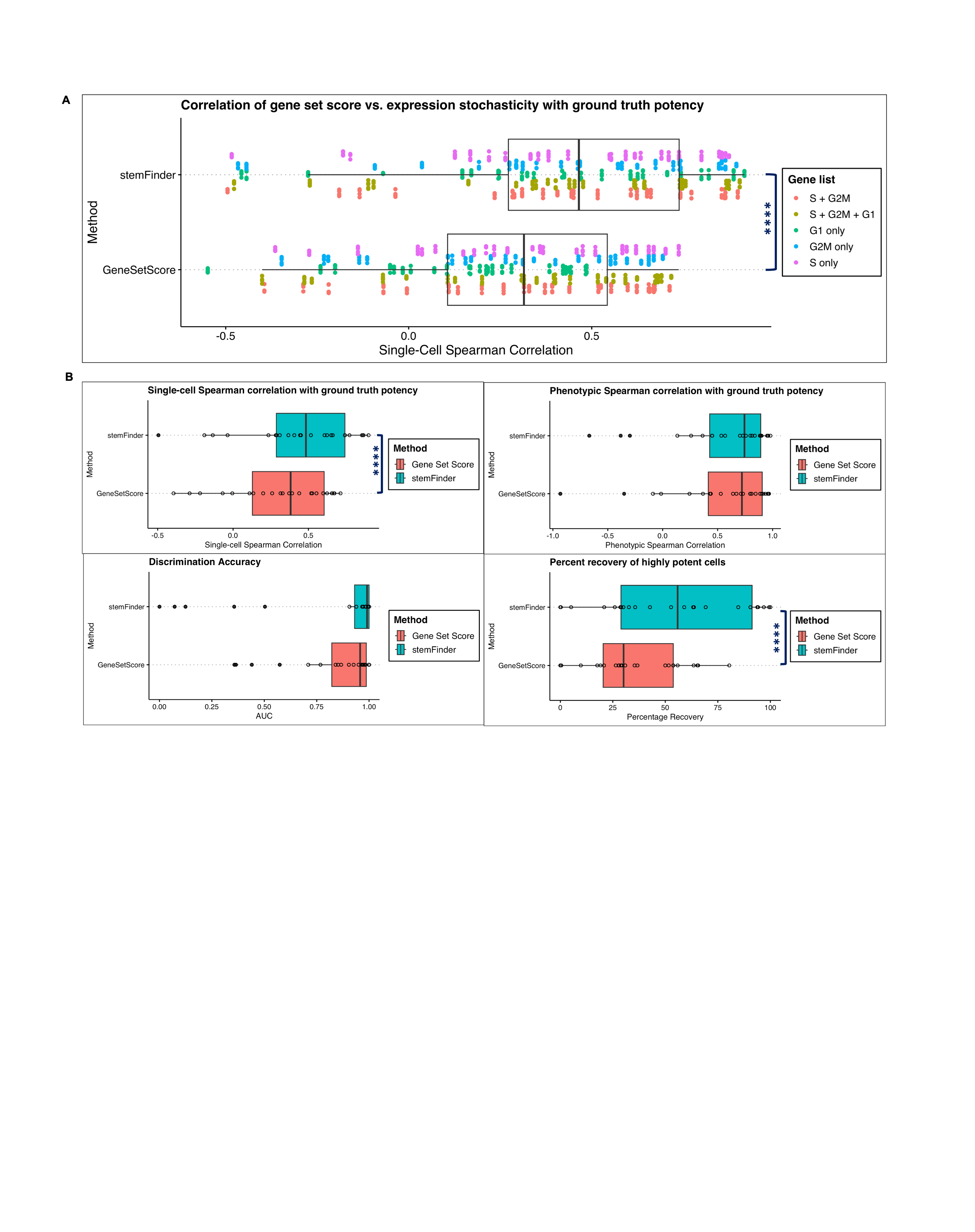

### FigS2AtoB.tiff

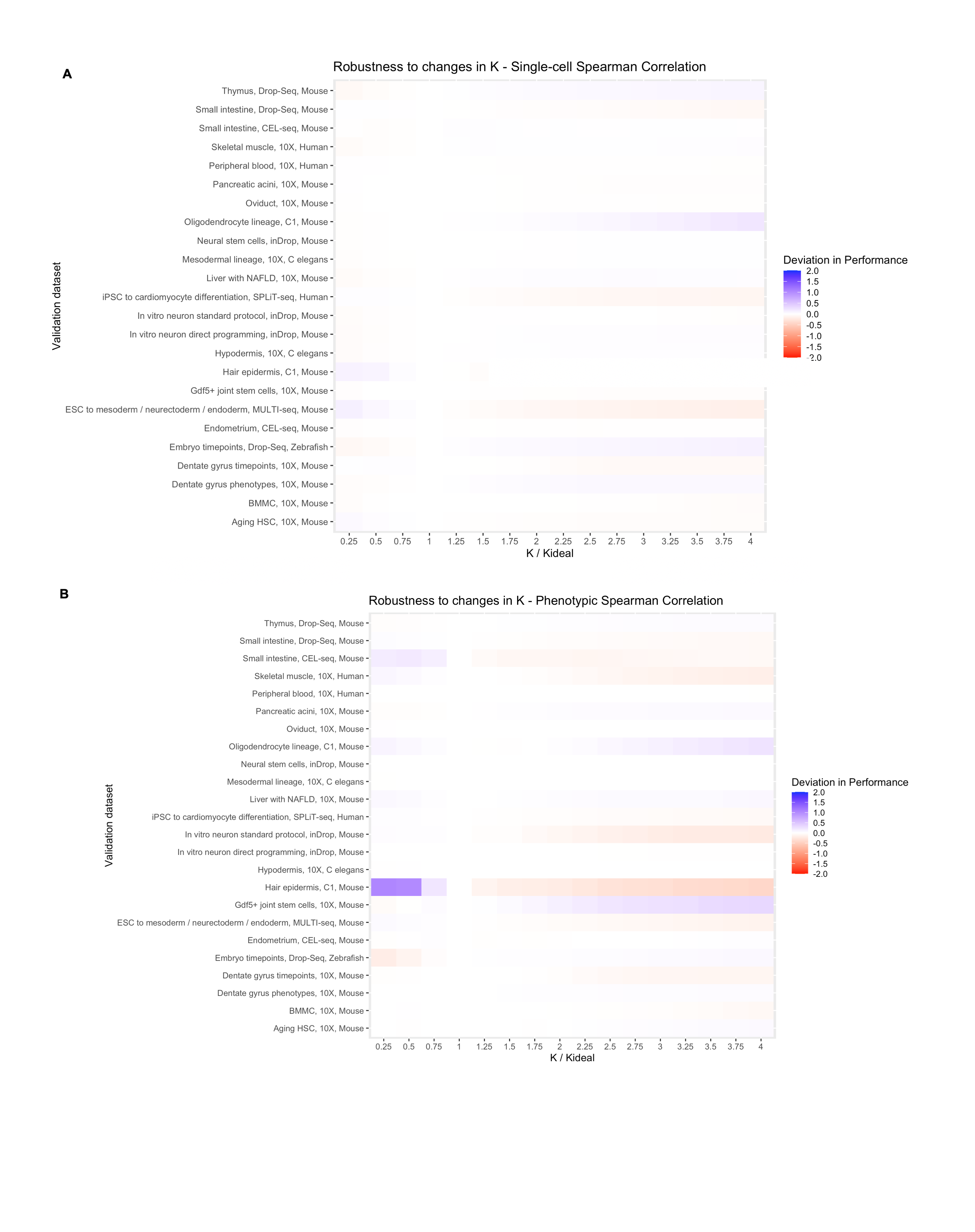

### FigS2CtoD.tiff

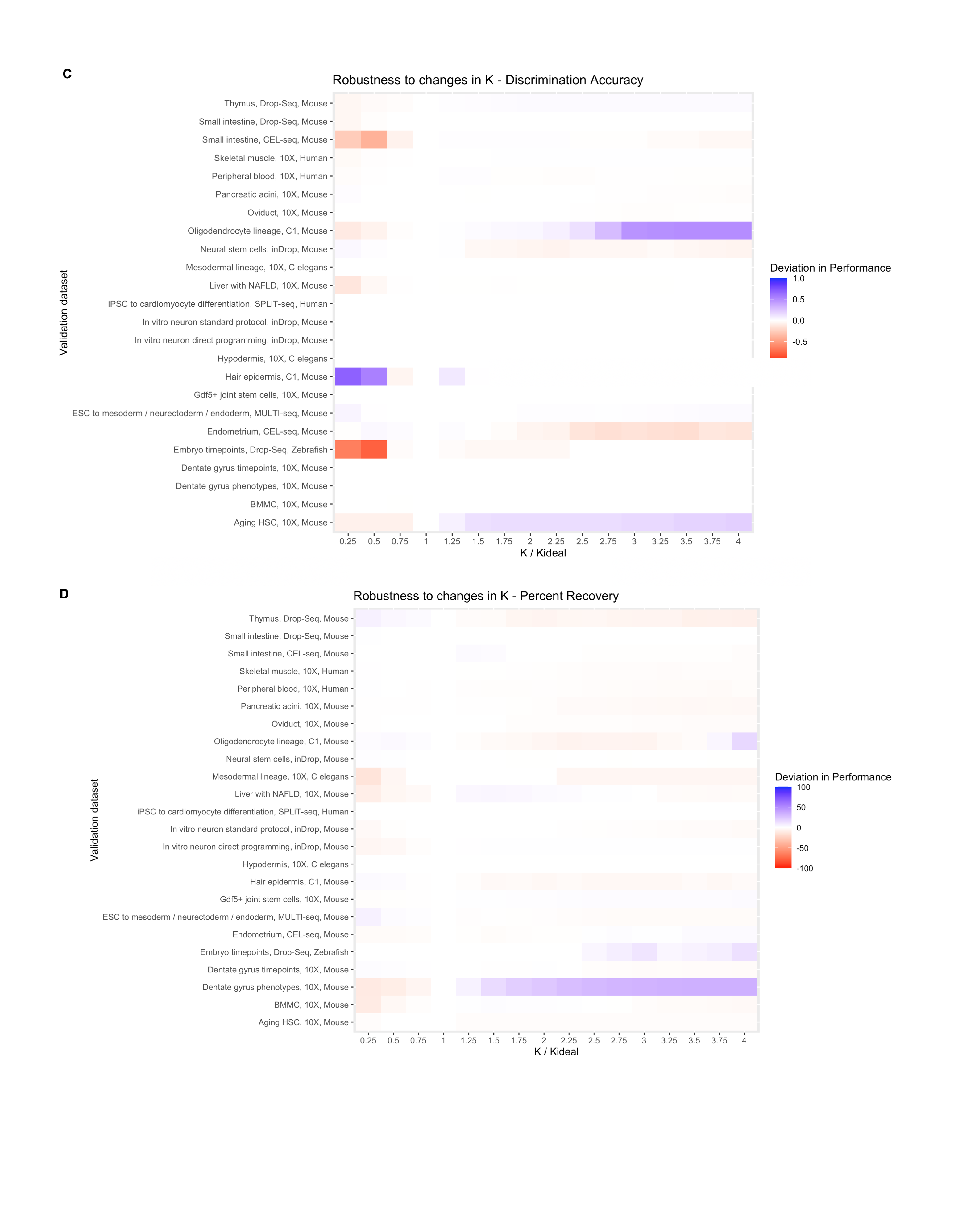

### FigS2EtoF.tiff

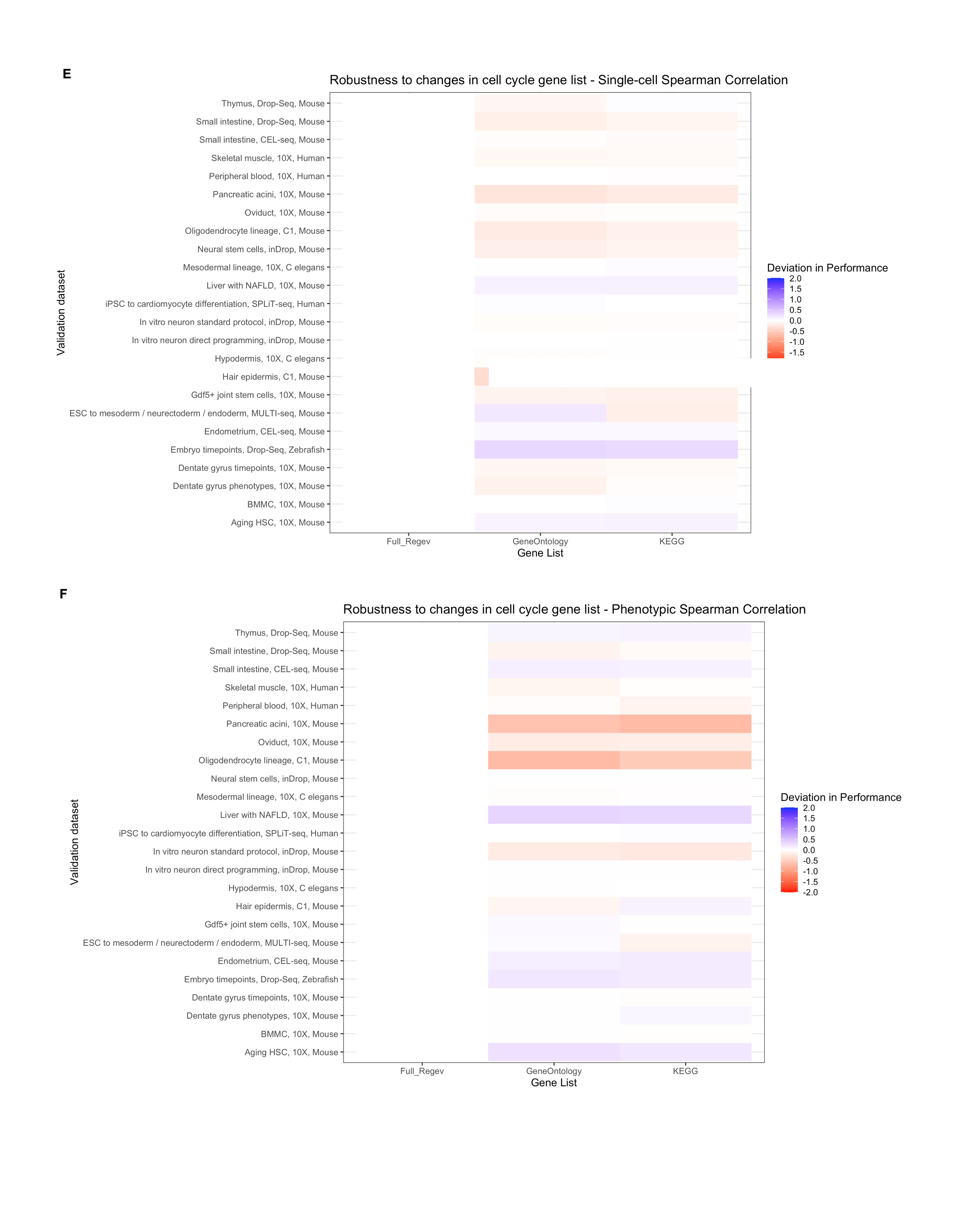

### FigS2GtoH.tiff

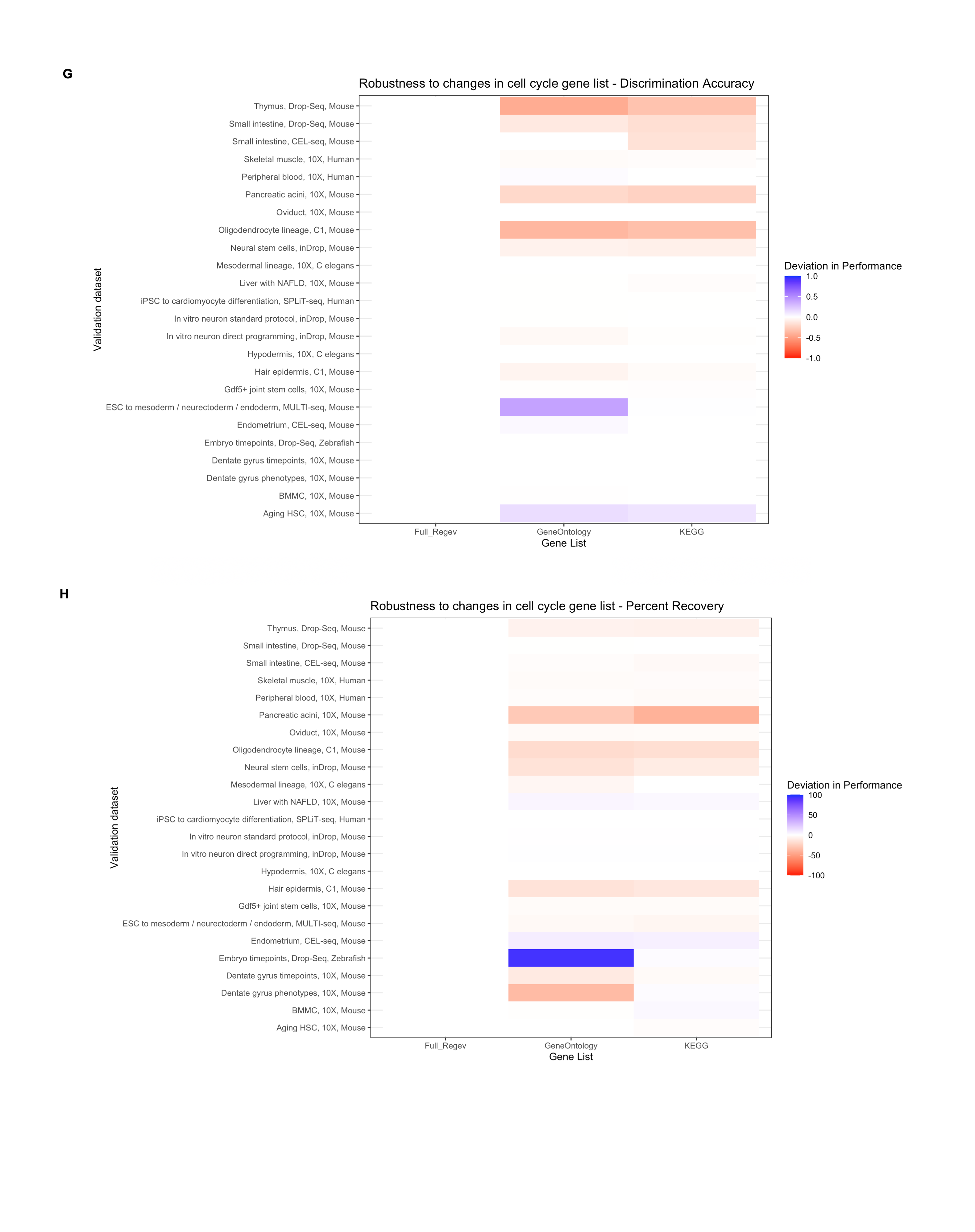

### FigS2ItoJ.tiff

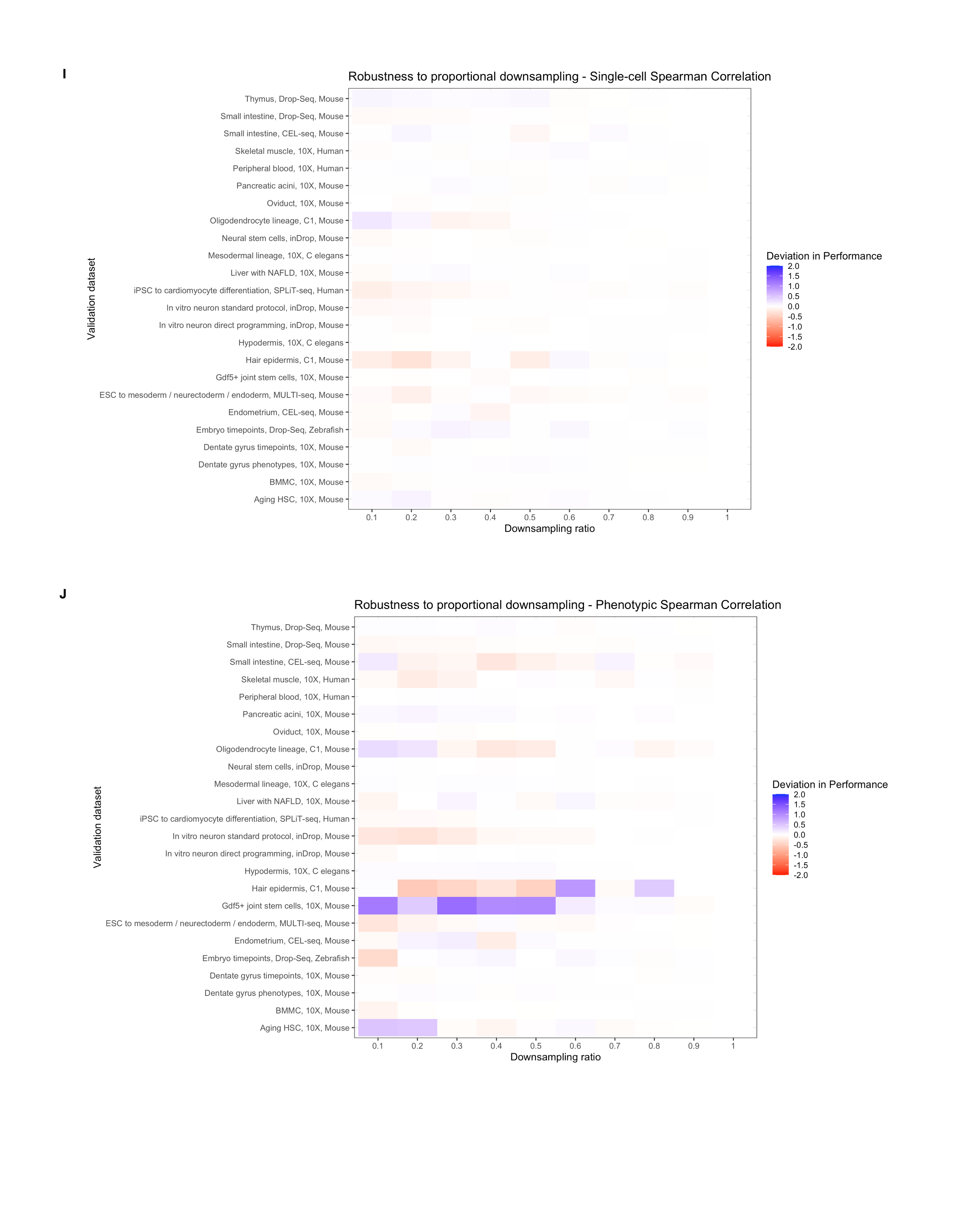

### FigS2KtoL.tiff

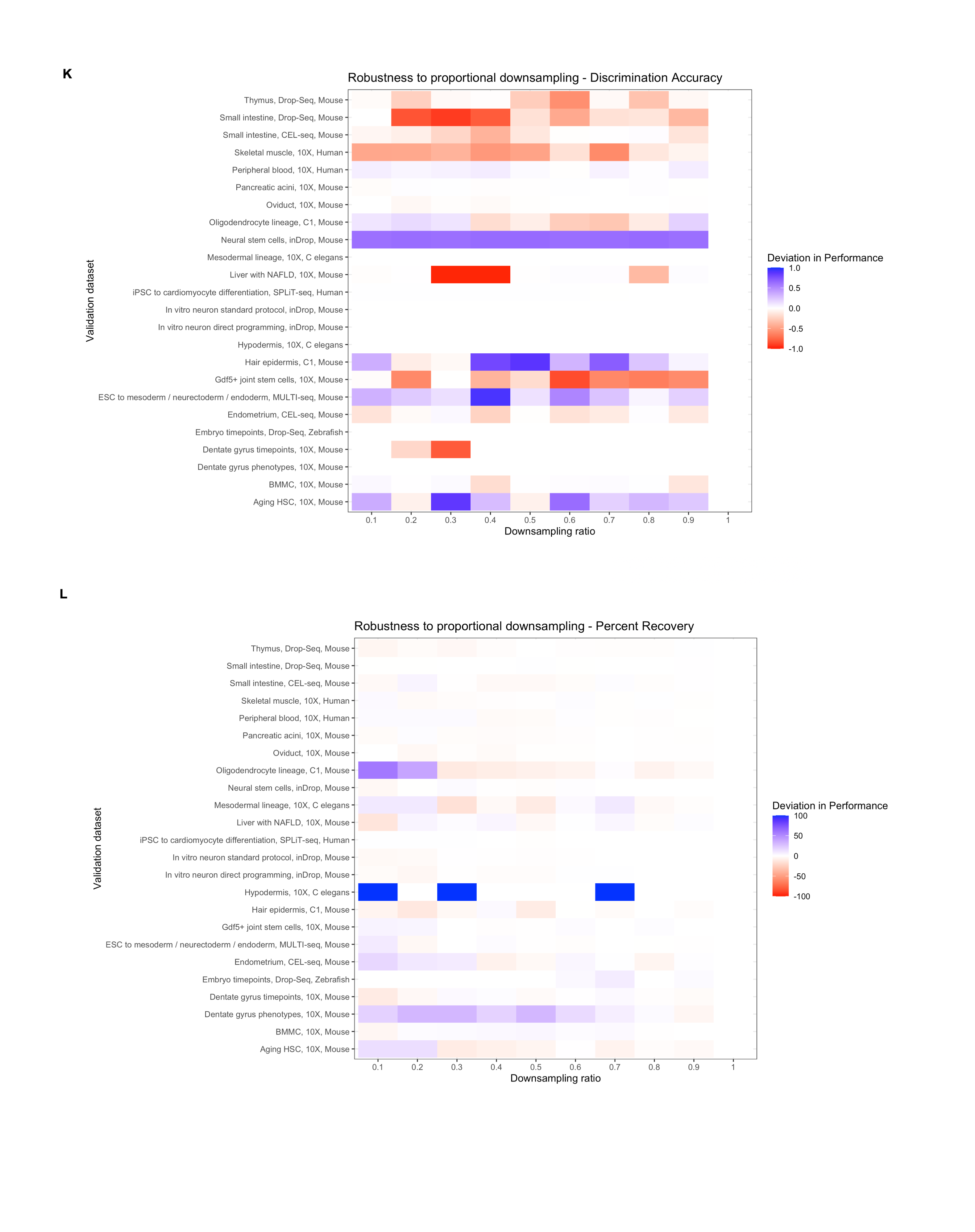

### FigS2MtoN.tiff

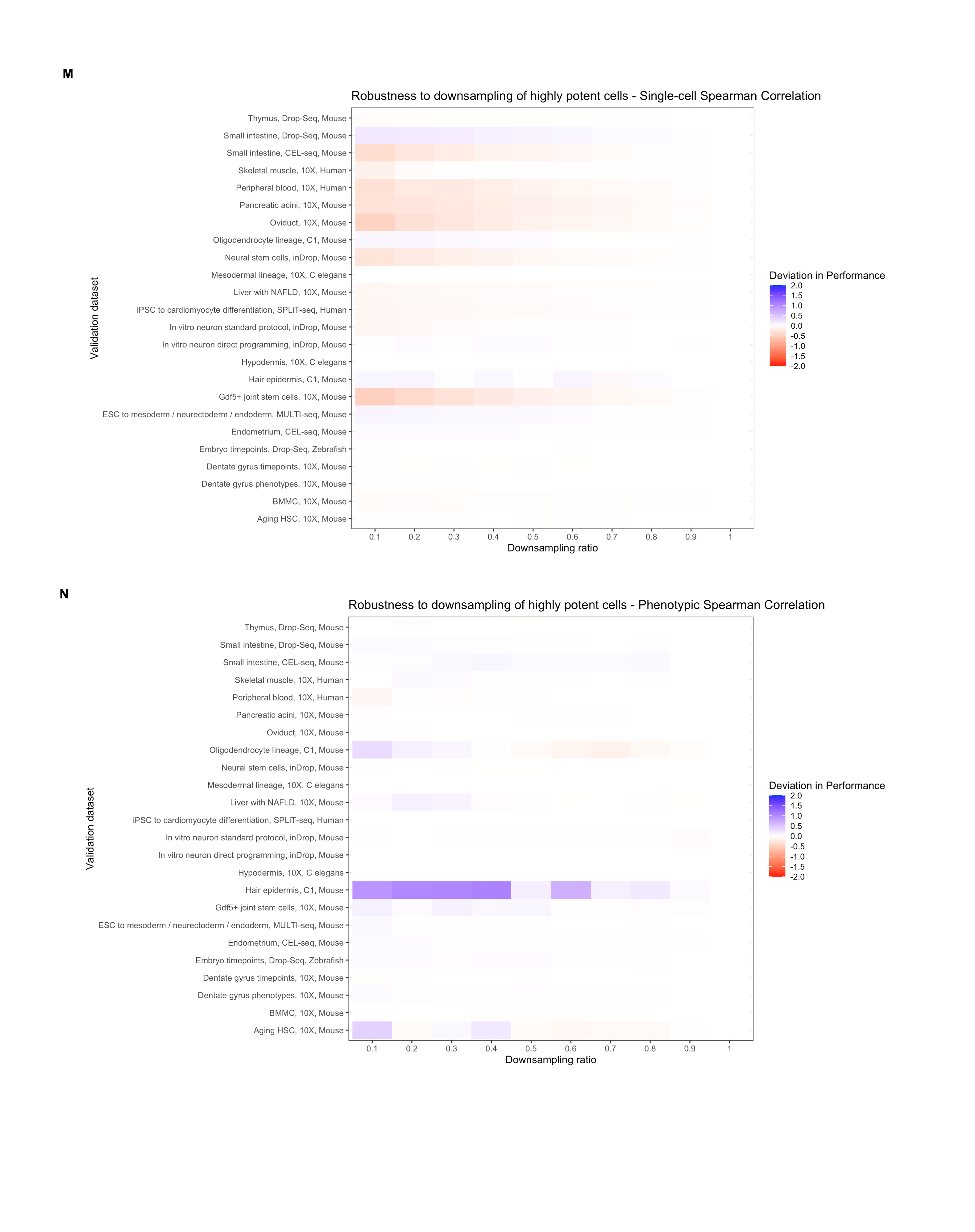

### FigS2OtoP.tiff

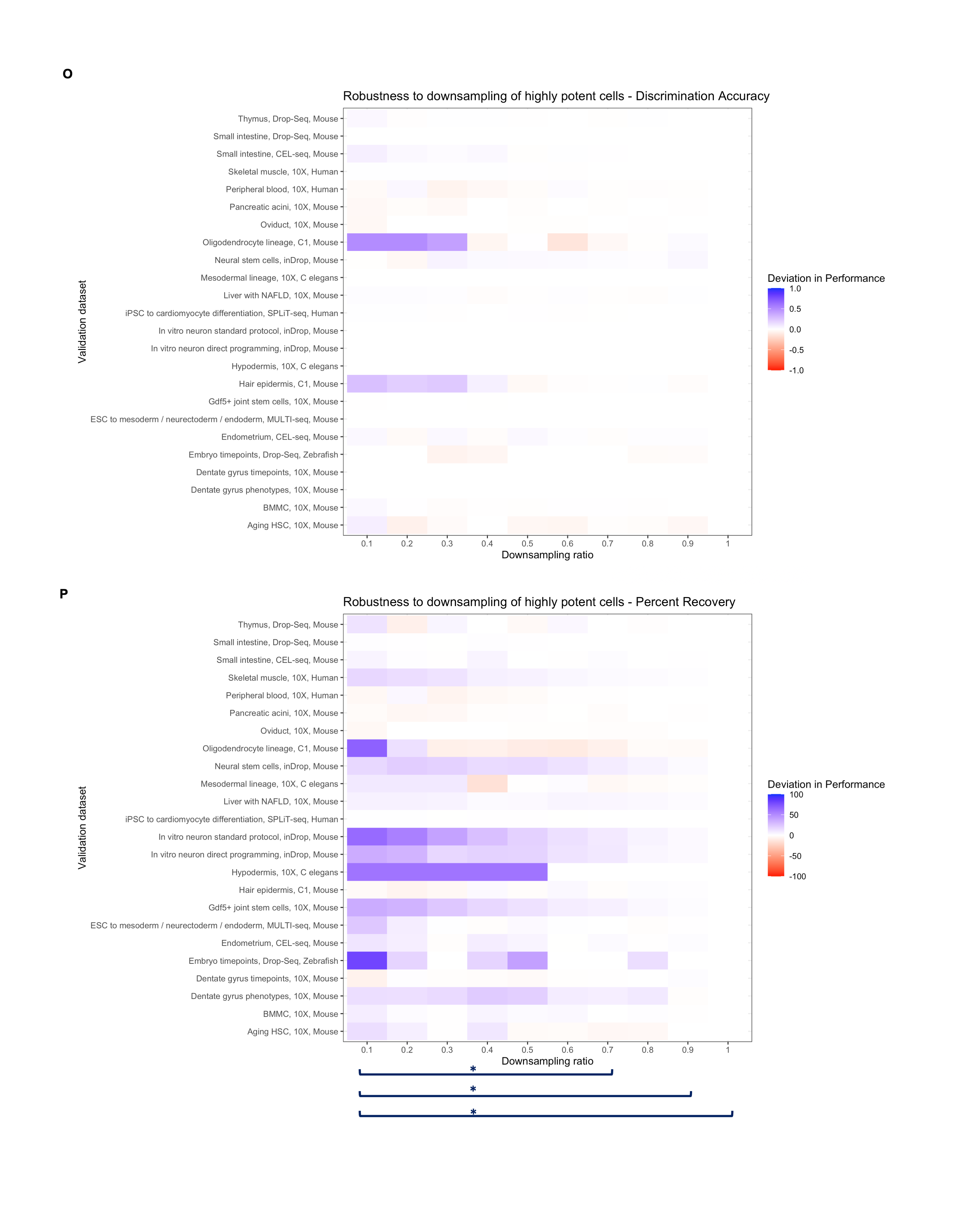

### FigS2QtoT.tiff

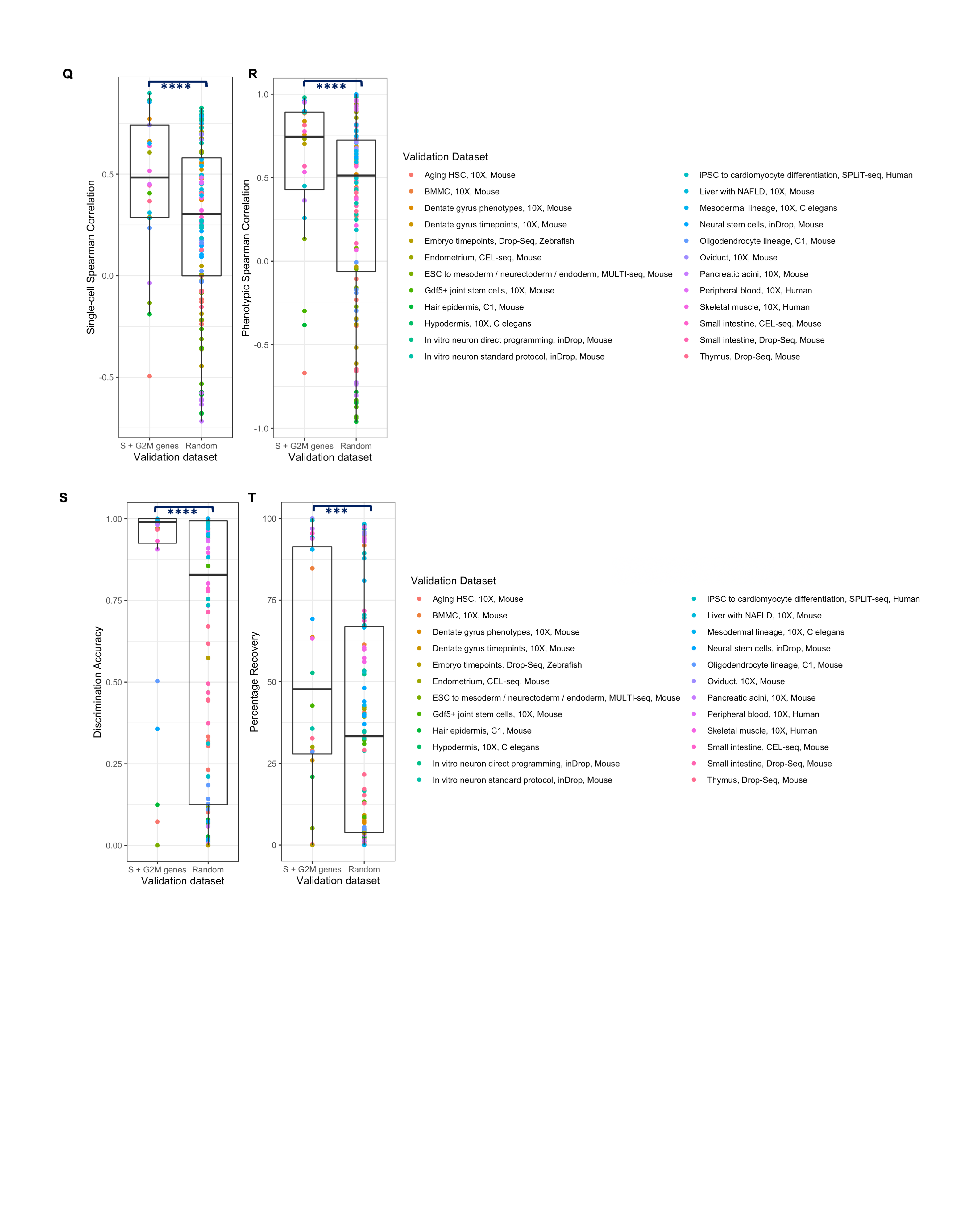

### FigS3.tiff

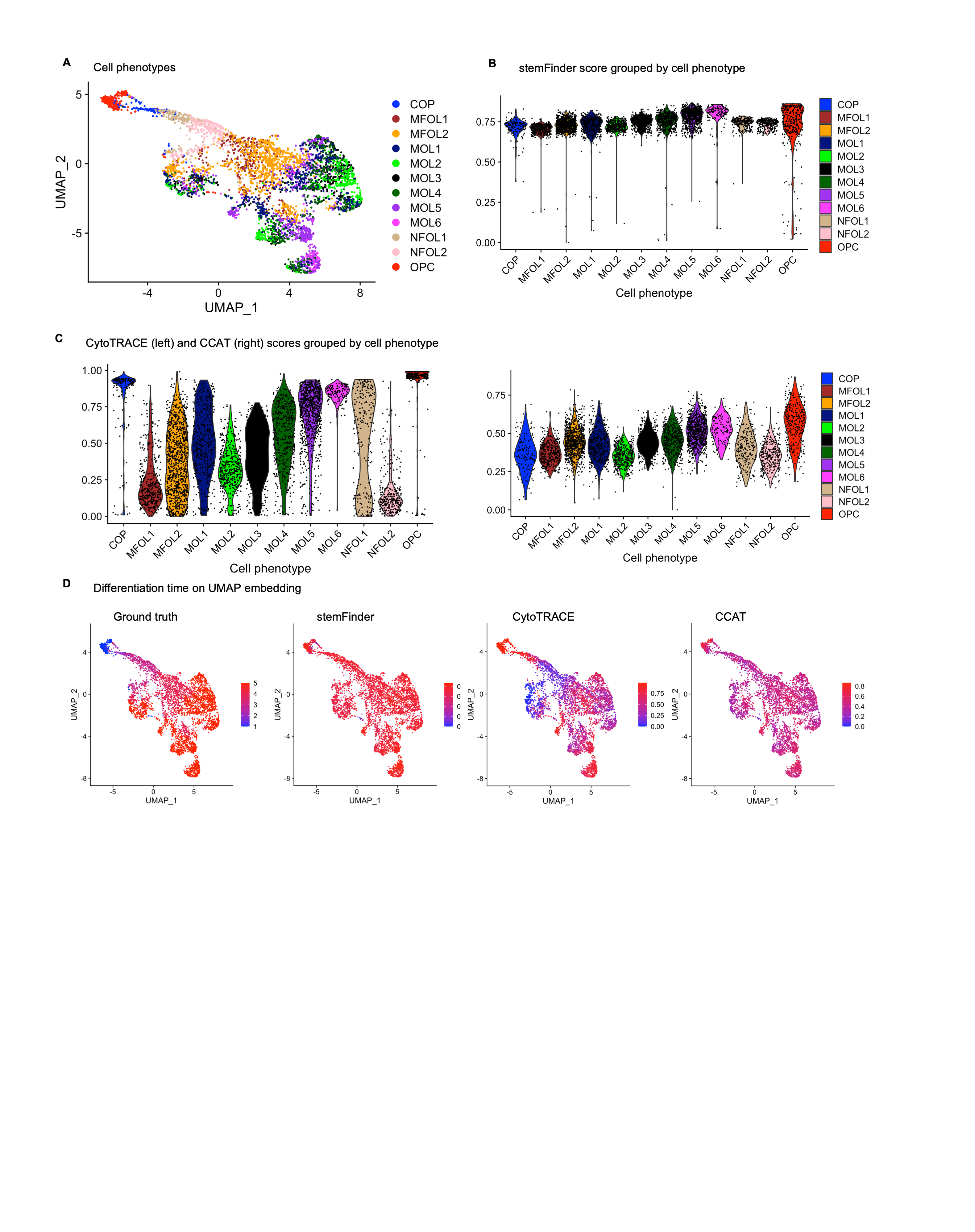
